# Transcriptional control of *hgcAB* by an ArsR*-*like regulator in *Pseudodesulfovibrio mercurii* ND132

**DOI:** 10.1101/2022.10.17.512643

**Authors:** Caitlin M. Gionfriddo, Ally Bullock Soren, Ann Wymore, D. Sean Hartnett, Mircea Podar, Jerry M. Parks, Dwayne A. Elias, Cynthia C. Gilmour

## Abstract

The *hgcAB* gene pair encodes mercury (Hg) methylation capability in a diverse group of microorganisms, but its evolution and transcriptional regulation remain unknown. Working from the possibility that the evolutionary function of HgcAB may not be Hg methylation, we test a possible link to arsenic resistance. Using model Hg-methylator *Pseudodesulfovibrio mercurii* ND132, we specifically evaluated transcriptional control of *hgcAB* by a putative ArsR encoded upstream and co-transcribed with *hgcAB*. This regulator shares homology with ArsR repressors of arsenic resistance and S-adenosyl-homocysteine (SAH) responsive regulators of methionine biosynthesis but is distinct from other ArsR/SahR in *Pseudodesulfovibrio mercurii* ND132. Using qPCR and RNA-seq analyses we confirmed this ArsR regulates *hgcAB* transcription, and is responsive to arsenic and SAH. Additionally, RNA-seq indicated a possible link between *hgcAB* activity and arsenic transformations by *Pseudodesulfovibrio mercurii* ND132, with significant up-regulation of other ArsR-regulated arsenic resistance operons alongside *hgcAB*. Interestingly, wild-type ND132 was less sensitive to AsV (but not AsIII) than an *hgcAB* knockout strain, supporting the idea that *hgcAB* may be linked to arsenic resistance. Arsenic significantly impacted Hg-methylation rates by ND132, however, responses varied with culture conditions. Differences in growth and overall metabolic activity did not account for arsenic impacts on methylation. One goal of this research is to better predict MeHg production in nature. However, we found that *hgcAB* gene and transcript abundance was not a good predictor of Hg-methylation rates. Our finding that *hgcAB* activity is linked to arsenic may hold clues to the possible environmental drivers of horizontal transfer of *hgcAB*.

**IMPORTANCE:** This work reveals a link between microbial mercury methylation and arsenic resistance and may hold clues to the evolution of mercury methylation genes (*hgcAB*). Microbes with *hgcAB* produce methylmercury, a strong neurotoxin that readily accumulates in the food web. This study addresses a critical gap in our understanding about the environmental factors that control *hgcAB* expression. We show that *hgcAB* expression is controlled by an ArsR-like regulator responsive to both arsenic and S-adenosyl-homocysteine in our model organism, *Pseudodesulfovibrio mercurii* ND132. Exposure to arsenic also significantly impacted *Pseudodesulfovibrio mercurii* ND132 mercury methylation rates. However, expression of *hgcAB* was not always a good predictor of Hg methylation rates, highlighting the roles of Hg bioavailability and other biochemical mechanisms in methylmercury production. This study improves our understanding of the controls on *hgcAB* expression which is needed to better predict environmental methylmercury production.

## INTRODUCTION

Microorganisms play a major role in controlling the form and fate of mercury (Hg) in aquatic environments. Methylmercury (MeHg), an organic Hg compound, is a neurotoxin that accumulates in food sources and is primarily produced by microorganisms(1). The *hgcAB* gene pair that encodes Hg methylation capability(2) is widespread in anaerobic environments and present in genomes spanning a diverse group of metabolic clades(3–5). Hg methylation capability is a species-specific trait that is often associated with low abundant taxa in the microbiome(3, 6). Currently, we do not have a good metric for estimating the prevalence of the mercury methylation trait among microbiota. However, in environmental metagenomic studies of mercury methylator environments, sequencing coverage of *hgcAB* genes is often low compared to other functional genes(7). For example, the coverage of dissimilatory sulfite reductase genes (*dsr*) in metagenomic assemblies is often 3-34x higher than *hgcAB*(7). Despite the association of sulfate-reducing bacteria (SRB) with Hg methylation, the *hgcAB* gene pair is found in only ∼12% of publicly sequenced SRB in the Deltaproteobacteria (although *hgcAB* is more common within some genera including *Pseudodesulfovibrio*)(8–10). Of the roughly 3,078 sequenced Deltaproteobacteria genomes in the NCBI reference database, only 368 were identified as potential Hg-methylators when the Hg-MATE database was compiled(7, 11). Within other phyla, especially environmental metagenome assembled genomes from the *Bacteroidetes*, *Nitrospirae*, PVC superphylum and candidate phyla radiation, *hgcAB* distribution further diverges from species trees but follows environmental patterns and suggests horizontal gene transfer of this capability(4, 6). Not all Hg-methylators share the same methylation capabilities, indicating that Hg uptake, Hg(II) binding to HgcAB, *hgcAB* transcription, enzyme regulation, or protein structure may control Hg methylation capacity and efficiency(12–17). Currently, we have a limited understanding of the biochemical constraints on Hg-methylation and the environmental conditions that resulted in the conservation of *hgcAB* in Bacterial and Archaeal genomes.

Despite the identification of two genes that encode Hg methylation capability, the mechanism by which methylation is performed within the cell is largely speculative. The genes encode a corrinoid (i.e., vitamin B_12_-dependent) protein (HgcA), and a 2[4Fe-4S] ferredoxin (HgcB), that could be used as a methyl carrier and electron donor in corrinoid cofactor reduction, respectively (2, 18). Both genes (*hgcA* and *hgcB*) are required for Hg methylation (2, 19). Corrinoid-dependent methyltransferases are utilized in a range of microbial metabolic pathways in the methylation of amino acids, proteins, DNA, and other intermediates(20, 21). In anaerobic microorganisms, corrinoid-dependent enzymes drive methionine and acetyl-CoA production (22, 23). Both pathways have been linked to Hg methylation, and the methyl transfer to Hg is thought to occur via a similar mechanism (18, 24–27). HgcA consists of a cytoplasmic corrinoid binding domain and a transmembrane domain. The cytoplasmic domain of HgcA shares homology with the C-terminal domain of the large subunit of the corrinoid iron-sulfur protein (CFeSP) utilized in methyl transfer in the Wood-Ljungdahl (i.e., reductive acetyl-CoA) pathway (2, 18).

Further, the native function(s) of HgcA and HgcB remaining unknown. Although HgcAB function has been linked to one-carbon metabolism(28), we don’t know if these proteins carry out any functions independent of Hg methylation. Redox transformations of metals and metalloids are often part of detoxification mechanisms or energy production. However, biomethylation of Hg does not provide any observed benefit to the cell, and Hg methylation capability is not induced by Hg(II)(28–30). Deletion of the *hgcAB* gene pair does not appear to hinder cell growth or produce any major phenotype in *Pseudodesulfovibrio mercurii* ND132 (2, 28). While some photosynthetic bacteria have been shown to use Hg(II) as an electron acceptor during photosynthetic and fermentative growth(31), Hg-methylating organisms are not able to use Hg species as growth substrates(30). Resolving the native biochemical function of HgcAB is needed to identify the metabolic conditions that mediate HgcAB activity and function. Understanding the biological and geochemical factors that influence MeHg production is necessary for predicting the impacts of contamination and remediation efforts on natural systems.

Building predictive models for net MeHg production in aquatic environments is a challenge(32). There are several microbial, physical, and geochemical parameters that influence mercury biogeochemistry in aquatic environments, including but not limited to Hg complexation, availability of terminal electron acceptors, shifts in redox, the composition or availability of organic matter, and the presence, abundance, and activity of Hg-cycling microorganisms(1, 33, 34). While the *hgcAB* gene pair can be used to identify potential Hg-methylators in a microbiome(6), to date, *hgcAB* gene abundance alone has rarely provided a useful indicator of environmental MeHg production(35–38). Measuring metabolic and *hgcAB* activity of Hg-methylators has been proposed to improve predictions(36, 39–42). However, it is unclear whether we can relate *hgcA* activity to Hg methylation rates(29). There are several significant unknowns, including whether *hgcAB* is transcriptionally regulated, what geochemical conditions influence *hgcAB* transcription, and if higher expression of *hgcAB* transcripts increases MeHg production. Previous work indicates that *hgcAB* transcription is not regulated by Hg(28–30). A possible overlap with microbial mechanisms for arsenic resistance has been proposed, as several studies have shown that *hgcAB* is co-localized with genes that encode for arsenic resistance in some Hg-methylators(4, 8, 43), including an ArsR-like regulator. In Hg-methylating members of the *Pseudodesulfovibrio*, this *arsR*-like gene is often found with *hgcAB*, and has been shown to be co-transcribed with *hgcAB*(8).

In this study, we test the ability of the ArsR-like regulator co-transcribed with *hgcAB* to enhance both *hgcA* transcription and Hg methylation itself in *Pseudodesulfovibrio mercurii* ND132, a model Hg methylator(9, 30). Two transcriptional responses were tested, one to arsenic using arsenite (As(III)) and arsenate (As(V)), and the second to S-adenosyl-homocysteine (SAH) using methionine and adenosine-2’,3’-dialdehyde. We chose these test conditions due to the shared homology between this potential *hgcAB* regulator and helix-turn-helix regulons in other sulfate-reducing bacteria that are responsive to As(III) (ArsR) and S-adenosyl-homocysteine (SahR). These regulate arsenic resistance or methionine biosynthesis, respectively, a(44–46). There are possible biochemical overlaps between Hg-methylation and S-adenosyl-methionine (SAM)-mediated methylation(47, 48), including those utilized in methyl-arsenic or methionine biosynthesis. Experiments were designed to test whether *hgcAB* transcription is regulated by ArsR/SahR.

In addition to evaluating potential *hgcA* regulators, another goal of this study was to test the idea that Hg-methylation rates are related to *hgcAB* expression rates. We hypothesized that geochemical culture conditions that increase *hgcAB* expression in *Pseudodesulfovibrio mercurii* ND132 will lead to higher Hg methylation rates than analogous conditions. Outcomes of this study have important implications for modelling Hg-methylation potential in the environment.

## METHODS

### Bacterial strains and culture media

For this study, we used the sulfate-reducing Hg-methylator and incomplete-oxidizer *Pseudodesulfovibrio mercurii* ND132 (previously *Desulfovibrio desulfuricans* ND132) as a model organism for understanding the mechanism and genetic regulation of Hg methylation(9, 30). *P. mercurii* ND132 has a well annotated genome, can grow on a number of substrates including lactate/sulfate, pyruvate/fumarate, formate/acetate/sulfate, and formate/acetate/fumarate(49). Furthermore, knock-out strains (Δ*hgcA,B*) of *P. mercurii* ND132 have been used to show the requirement of *hgcA* and *hgcB* for Hg methylation(2, 19). For this study, we used *P. mercurii* ND132 wild-type and mutant glycerol stock cultures from Dr. Dwayne Elias’ culture collection. Since these experiments began, wild-type *Pseudodesulfovibrio mercurii* ND132 has been deposited into DSMZ (type strain DSM 110689) and ATCC (ATCC TSD-224) culture collections. The Hg-methylation knock-out strain (Δ*hgcAB*) of *P. mercurii* ND132 was originally sourced from Judy Wall(2). Stock cultures were stored at −80°C in lactate-sulfate media and 10% v/v glycerol.

ND132 is capable of fumarate respiration using pyruvate as an electron donor(9). For most assays, *P. mercurii* ND132 was grown in this way on either CCM(50) or Estuarine Pyruvate Fumarate (EPF) medium(9). Both media contain 40 mM pyruvate and 40 mM fumarate. CCM is a defined medium, while EPF includes 0.5% yeast extract. Both media were buffered to pH 7.2 using MOPS and include vitamin and mineral additions. Media were reduced with 0.1 mM (CCM) or 0.05 mM (EPF) cysteine-HCl to remove trace oxygen contamination. Other amendments varied by experiment. All media contained resazurin as a redox indicator. For some arsenic toxicity assays, ND132 was also grown on CCM media with lactate (60 mM) and sulfate (30 mM) as electron donor and acceptor. All culture work was done at 31 ± 1 °C.

### Experimental assays for testing regulators of Hg methylation and *hgcAB* transcription

We evaluated the potential transcriptional response of a putative *hgcAB* regulator to S-adenosyl-homocysteine (SAH) and arsenic. We also evaluated potential influence of *hgcA* expression on Hg methylation using batch culture and washed cell approaches. For all assays, cultures were started from freezer stocks. In all cases, cells were grown on CCM or EPF media. Out of the freezer, cells were grown to mid-late exponential phase, then inoculated at 10% v/v into medium containing one of the four test conditions, or unamended control medium. In defined CCM media the test amendments were: 0.2 mM sodium arsenite, 2.0 mM sodium arsenate, 1 mM methionine, 0.1 mM adenosine-2’3’-dialdehyde, and 1 nM of HgCl. In EPF media, arsenic concentrations were reduced to 0.02 mM sodium arsenite and 0.2 mM sodium arsenate. Amendment concentrations for expression and methylation studies were chosen based on toxicity tests (see below). Arsenic concentrations selected were the highest levels that allowed unimpacted growth, which was specific to each medium. In qPCR expression experiments conducted over the growth curve, both control and amendment cultures were grown without added Hg. In the washed-cell assays and Hg-methylation assays performed in culture, all treatments including the control were spiked with 1 nM isotopically enriched ^201^Hg.

Methylation assays were carried out in triplicate by the addition of 1 nM isotopically enriched ^201^Hg (98.1% purity; Oak Ridge National Laboratory) (in 1% v/v HCl). Uninoculated sterile medium with and without treatment amendments served as abiotic controls. Methylation assays ran 100 minutes, after which aliquots of unfiltered culture were immediately acidified (0.5% v/v HCl) and frozen (−20°C) for Me^201^Hg analysis. Experimental timeframes were selected on the basis of previous kinetic experiments demonstrating a plateau in MeHg production by strain ND132(17).

Methylation assays were performed using either washed cells or batch culture cells. For washed cell assays, cells were harvested at mid log phase (OD_600_ ∼ 0.2) from batch culture and washed and resuspended in a minimal wash buffer as described previously(51). The salt buffer contained 1 mM pyruvate and 1 mM fumarate, reduced with 0.5 mM cysteine-HCl, and buffered to pH 7.5. For batch culture experiments, culture aliquots from mid log phase (OD_600_ ∼ 0.2) were directly spiked with ^201^Hg. In most cases, the concentration of filterable and total ^201^Hg was measured immediately after the ^201^Hg spike, and at the end of the methylation assays (and in some cases through time during the assays). Samples were filtered through 13mm diameter 0.22 μm CellTreat® MCE membrane syringe filters. For most studies we also measured pH, optical density at 600 nm wavelength (OD_600_), acridine orange direct cell counts (AODC), and sulfide concentration over the course of the assay. Additionally, for some studies we measured short-chain fatty acids and/or CO_2_ production as indicators of cellular activity. All culture manipulations were carried out in a Coy® glove bag under an O_2_-free atmosphere.

Culture aliquots were saved for RNA extraction before the assays and from each assay replicate at the end of the time-course (t = 100 minutes). For RNA extraction, cells were harvested from batch cultures and methylation assays at 2,800 x g for 10 minutes. The cell pellet was resuspended in 1 mL TRIzol™ Reagent (Invitrogen), and kept at −20°C prior to nucleic acid extraction.

### Hg and MeHg analyses

Isotope-specific total Hg and MeHg concentrations were measured using modifications of EPA Methods 1631 (EPA-821-R-01-023, Mar 2001) and 1630 (EPA-821-R-01-020, Jan 2001) respectively, with inductively coupled plasma mass spectrometry (ICP-MS) detection. Details of the analytical method and QA/QC (Table S1) are provided in the SI.

### Other analytical methods

Sulfide was analyzed using a sulfide selective electrode with an Ag−AgCl reference electrode (Thermo Scientific) and was calibrated against Pb-titrated Na_2_S standards made in sulfide antioxidant buffer (SAOB). Saturated Na_2_S stocks were made from washed crystals in degassed ultra-pure water and held in N_2_-filled sealed serum vials until use. SAOB contains 2 M NaOH, 0.2 M EDTA and 0.2 M ascorbic acid, in degassed ultra-pure water and sealed under N_2_ until use. Organic acids (acetate, pyruvate, fumarate, and succinate) (0.5‒ 200 μM) were determined with a Dionex^TM^ Integrion HPIC equipped with an AS11-HC column at 35 °C with a KOH gradient of 1‒60 mM with an ICS-5000^+^ EG effluent generator system (Thermo Fisher Scientific). Sample volumes of 10 μl were injected with a Dionex^TM^ AS-AP auto-sampler at room temperature.

Cell counts were done manually using acridine orange staining. Cells were fixed at a dilution of 1:100, with 0.1 mL of cell culture preserved in 9.1 mL of 20 mM phosphate buffer solution (PBS) and 0.8 mL glutaraldehyde, and stored in the fridge prior to slide prep. Slides were made by combining 0.2 mL of 0.1% acridine orange and 1.9 mL of fixed cells in microcentrifuge tubes and allowing 1 hr at room temperature in the dark for staining. The stained cells were mixed with 1 mL of 20 mM PBS and collected on filters via vacuum. Counts were performed manually and averaged from 10 different grids over a 0.01 mm^2^ area.

CO_2_ production was measured over time in the headspace of serum vials containing culture aliquots, after purging headspace w/ O_2_-free N_2_ for ∼5 min. All measurements were made from duplicate aliquots for each treatment. The 100 ml serum vials contained either 5 ml of culture, or 25 mL of washed cells in salt buffer. Headspace CO_2_ was measured hourly after an initial equilibration period using a LICOR LI-7000 CO_2_ analyzer. Total inorganic carbon production was calculated from headspace CO_2_ concentrations based on temperature, pH, the Henry’s Law constant for CO_2_ and equilibrium among dissolved inorganic carbon (DIC) species. DIC production rates were calculated from the linear portion of the slope with time

### RNA extraction

Cell cultures were pelleted at 2,800x g for 10 minutes, excess media removed, and then resuspended in 1 mL of TRIzol^TM^ reagent (Invitrogen), and stored at −20 °C until processing. RNA was extracted following TRIzol protocol, which involves a phenol-chloroform extraction, isopropanol precipitation, and ethanol cleaning step. Extracted RNA was further purified using RNeasy Clean-up kit (Qiagen), and DNAse treatment with TURBO DNA-free™ Kit (Thermo Fisher). RNA concentration was measured using a Qubit™ RNA HS assay kit on the Qubit 3.0 Fluorometer (Thermo Fisher), and purity was determined by NanoDrop (Thermo Fisher). gDNA contamination was assessed by performing a qPCR with DNase-treated RNA prior to reverse transcription. cDNA synthesis was performed using iScript™ cDNA Synthesis Kit (Biorad), and cDNA concentration was measured using Qubit™ ssDNA assay kit on the Qubit 3.0 Fluorometer (Thermo Fisher). cDNA was diluted to 1 ng/uL with RNase-free water prior to qPCR.

### Real-time quantitative PCR (qPCR)

qPCR reactions were set up on a 96-well plate with each standard and biological cDNA sample run in triplicate. Each well contained 1 ng of cDNA, 0.25 µM forward and reverse primer, 0.02 µL ROX reference dye (Bio-Rad), 10 µL iQ SYBR Green Supermix (Bio-Rad), and 1xTE Buffer (pH 8), filled to 20 µL total volume. For standard curves, we used serial dilutions of 4.2 ng/µL (measured on Qubit™) gDNA extracted from pure culture *P. mercurii* ND132 wild type, estimated to be equivalent to 10^6^ copies/µL based on the size of the genome (3.8 Mbp)(49). All standards and samples of each primer set were amplified on the same plate, with each plate only containing one target or reference gene. All qPCR analyses from the same experiment were performed in succession with the same standard curve. Custom-designed oligo primers were used to amplify 107 nt bp target region of *hgcA* (DND132_1056-hgcA-F, DND132_1056-hgcA-R) and 180 nt bp region of the *arsR*-like gene before *hgcA* (DND132_1054-arsR-F, DND132_1054-arsR-R) (Table S2). Three single-copy housekeeping genes that encode recombinase subunit A (*recA*), gyrase subunit A (*gyrA*), and gyrase subunit B (*gyrB*) were used for normalization of expression of genes of interest. These housekeeping genes were chosen for normalization due to their observed expression stability in ND132 wild-type transcriptomes (data not shown). Custom primers were designed to amplify a 153 nt bp region of *recA* (DND132_1603, DND132_1603-recA-F, DND132_1603-recA-R), 148 nt bp region of *gyrA* (ND132_2285, DND132_2285-gyrA-F, DND132_2285-gyrA-R), and 182 nt bp of *gyrB* (DND132_2284, DND132_2284-gyrB-F, DND132_2284-gyrB-R). Primers were designed to function over the same annealing temperature (65 °C), and the same thermocycler protocol was used to amplify each gene: polymerase activation for 3 minutes at 95 °C, followed by 32 cycles of denaturing for 15 seconds at 95 °C, and annealing for 20 seconds at 65 °C, followed by a melt curve of 0.1 °C/s from 65 °C to 95 °C, ending with 15 seconds at 95 °C.

### RNA-seq transcriptomics

RNA sequencing was performed on clean, DNase-treated RNA extracted from the batch culture and washed-cell Hg-methylation assays. For the washed-cell assays, only RNA from the control, As(III), and As(V) treatments were sent for RNA-seq. All treatments from the culture Hg methylation assays were sent for sequencing. Three biological replicates for each treatment were sequenced. Library prep and sequencing were performed by Genewiz (Azenta Life Sciences, NJ). Ribosomal RNA (rRNA) depletion was performed during library preparation using three probes from QIAGEN FastSelect rRNA 5S/16S/23S Kit (Qiagen, Hilden, Germany), respectively. RNA sequencing libraries were prepared using NEBNext Ultra II RNA Library Preparation Kit for Illumina by following the manufacturer’s recommendations (NEB, Ipswich, MA, USA). Briefly, enriched RNAs are fragmented for 15 minutes at 94 °C, then first and second strand cDNA are synthesized. cDNA fragments are end repaired and adenylated at the 3’ ends, and universal adapters are ligated to cDNA fragments, followed by index addition and library enrichment with limited cycle PCR. Sequencing libraries were validated using the Agilent Tapestation 4200 (Agilent Technologies, Palo Alto, CA, USA), and quantified using Qubit 2.0 Fluorometer (ThermoFisher Scientific, Waltham, MA, USA) as well as by quantitative PCR (KAPA Biosystems, Wilmington, MA, USA). Multiplexed and clustered sequencing libraries were sequenced using Illumina Hiseq 2×150 bp paired-end configuration.

Raw sequencing files were uploaded to KBase(52). Initial read quality was assessed with FastQC, followed by adapter and quality-based trimming with Trimmomatic(53), alignment to the *Pseudodesulfovibrio mercurii* ND132 reference genome (https://www.ncbi.nlm.nih.gov/assembly/GCF_000189295.2) with HISAT2(54), assembly of transcripts from alignment using StringTie(55). Differential expression analysis was performed with DeSeq2(56) and the results were uploaded to Galaxy web platform(57), and we used the public server at usegalaxy.org to produce figures using the Galaxy R scripts.

### Arsenic analyses

Arsenic speciation analyses were done at Brooks Applied Labs (Bothell, WA, USA). Samples were collected from culture bottles, filtered (Whatman® cellulose acetate membrane syringe filters 0.45 μm) into BD Vacutainers®, preserved in EDTA and stored at 4 °C in the dark until analyzed. Arsenic (As) speciation analysis, including arsenite [As(III)], arsenate [As(V)], monomethylarsonic acid [MMAs], dimethylarsinic acid [DMAs], and unknown As species were determined on all samples. Arsenic species were chromatographically separated on an ion exchange column and then quantified using inductively coupled plasma collision reaction cell mass spectrometry (ICP-CRC-MS). The results were not method blank corrected and were evaluated using reporting limits adjusted to account for sample aliquot size.

### Structural modeling

Structural models of the putative ArsR-like transcriptional regulator from *P. mercurii* ND132 (DND132_1054, Uniprot ID: F0JBE8) were generated with ColabFold(58). ColabFold is an implementation of AlphaFold2(59) that uses MMSeqs2(60) to generate large multiple sequence alignments by searching the UniRef and MGnify databases(61). Based on its sequence similarity to other transcriptional regulators that are known to function as homodimers, we modelled the homodimeric form of the protein using AlphaFold-multimer(62), which is a version of AlphaFold2 trained specifically to model protein complexes. Models were ranked by their average predicted local distance difference test (pLDDT) and predicted TM-score (pTMscore) and the model with the highest pTMscore score was selected for further analysis.

### Statistical methods

Statistical modelling was performed in JMP®, Version 16 (SAS Institute Inc., Cary, NC, 1989–2022) and R statistical software (version 4.2.0)(63). Generalized linear models were used to evaluate differences in Hg methylation rates, inorganic carbon production, organic acid production, and growth rates among treatments. Where appropriate, models included time, treatment and any potential interaction terms. In the absence of significant interactions, differences among treatments were evaluated using pairwise comparison of slopes. For qPCR results, a two-sample z-test was used to determine if the difference in *arsR* and *hgcA* expression was significant between treatments and control. For significance testing, the mean and standard deviation of ΔΔCq values for each target gene and treatment were compared to control.

## RESULTS AND DISCUSSION

### An ArsR-like regulator encoded with HgcAB in *Pseudodesulfovibrio*

A putative arsenic responsive regulator (ArsR) (Gene ID: DND132_1054, Uniprot: F0JBE8) is encoded directly upstream of Hg-methylation genes *hgcA* (Gene ID: DND132_1056, Uniprot: F0JBF0) and *hgcB* (Gene ID: DND132_1057, Uniprot: F0JBF1) in the genome of *P. mercurii* ND132. Potential ArsR-controlled transcriptional regulation of *hgcAB* has been proposed for Hg-methylating members of the *Pseudodesulfovibrio*(8). This potential *hgcAB* regulator shares homology to the metalloregulator ArsR/SmtB protein family (Pfam PF01022), including a DNA-binding helix-turn-helix (HTH) motif. This family of regulators (Pfam PF01022) are responsive to metal ion stress (e.g., As(III), Zn(II), Cd(II)) and in the absence of coordinating metal(loid)s will repress the expression of operons linked to heavy metal resistance(64). This transcriptional family includes the ArsR repressor of the *ars* operon, which often encodes an arsenate reductase with arsenic permeases and exporters(65). However, protein sequence analysis and predictive structure modelling of the *hgcAB* ArsR-like regulator suggests differences in the metal binding domain that may make it distinct from other ArsR family regulators.

There are five ArsR family transcriptional regulators annotated in the genome of *Pseudodesulfovibrio mercurii* ND132 (Figure 1). Based on protein sequence analysis, these five ArsR proteins are different from one another, with less than 33% pairwise identity between them (Table S3). We numbered the *arsR* by order of appearance in the genome of *P. mercurii* ND132, and *arsR2* is located directly upstream of *hgcA* and *hgcB* (Figure 1). While in some Hg-methylating bacteria, *hgcAB* is located in the genome with arsenic resistance (*ars*) genes(4, 43), no other *ars* genes are colocalized with *hgcAB* in ND132. However, three of the other *arsR* (*arsR1*, *arsR3*, and *arsR5*) are present with *ars* genes elsewhere in the genome and likely regulate arsenic resistance. The first set of *ars* genes encode ArsR1 (GeneID: DND132_0890) and the first of two methylarsenite efflux permeases, ArsP (DND132_0889)(66). Another *ars* operon encodes ArsR3 (DND132_1317) with the first of two Acr3 family As(III) permeases (DND132_1318) and arsenate reductase, ArsC (DND 132_1319). *P. mercurii* ND132 has a second Acr3 family As(III) efflux permease that is not part of an *ars* operon (DND132_2196). The third *ars* operon is predicted to encode ArsR5 (DND132_2311), another ArsP (DND132_2312), an organoarsenical resistance protein, ArsE (PFam: PF13192, thioredoxin-like protein) (DND132_2313)(67), and a protein of unknown function with a diguanylate cyclase (DGC) motif (Pfam: PF08859, a domain with four conserved cysteines that likely coordinate Zn) (DND132_2314). ArsR repressors of the *ars* operon are predicted to be transcriptionally regulated by As(III) and part of a detoxification pathway for arsenic. From protein sequence analysis of ND132 ArsR2, it is unclear whether expression of *hgcA* and *hgcB* is part of the ArsR regulon and controlled by arsenic.

**Figure 1.**
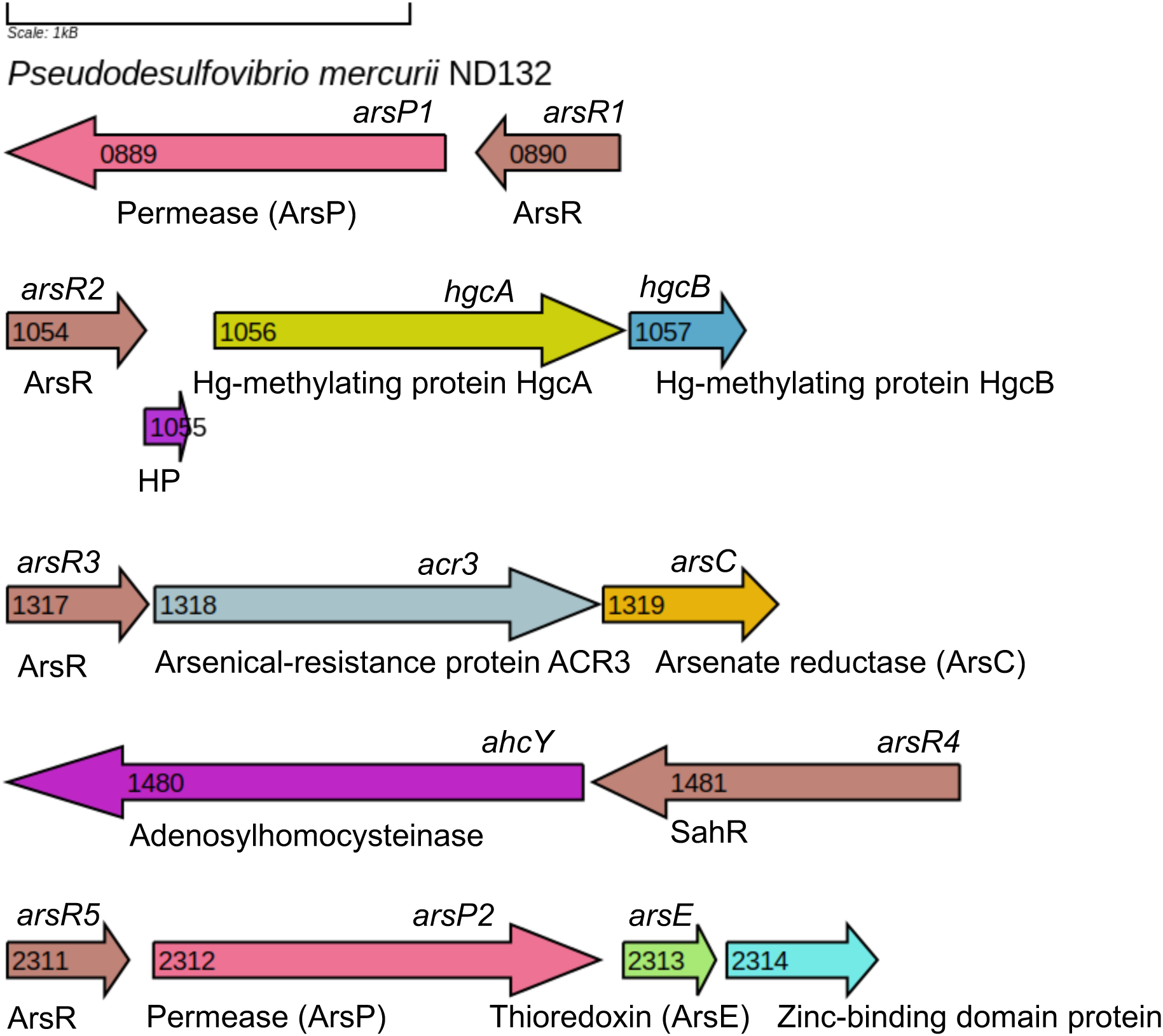
ArsR/SmtB family transcriptional regulators encoded in the genome of *Pseudodesulfovibrio mercurii* ND132. The five *arsR* genes are numbered by order of appearance in genome, with gene ID in arrow (e.g. HgcA enoded by DND132_1056), gene orientation shown by arrows, and color coded by protein family. Figure made using Gene Graphics(93).

ArsR/SmtB family regulons may not always be associated with heavy metal resistance or arsenic transformations. In addition to regulating arsenic resistance, ArsR regulators have also been shown to influence many other cellular functions in the presence of As(III), including phosphate acquisition/metabolism, sugar transport, chemotaxis, copper tolerance, and iron homeostasis(68). Regulators in the ArsR/SmtB HTH family may also be responsive to substrates other than heavy metals and metalloids. In sulfate-reducing bacteria, an ArsR-like regulon responsive to intracellular concentrations of S-adenosylhomocysteine (SAH) controls SAM-dependent methyl transfer for methionine biosynthesis(44, 45). These SAH-responsive regulators (SahR) are often encoded with adenosylhomocysteinase (AHCY), which catalyzes the hydrolysis of SAH to homocysteine and adenosine. In *P. mercurii* ND132, *arsR4* likely encodes a SahR (Gene ID: DND132_1481, Uniprot: F0JE80). The predicted SahR is encoded with AHCY (Gene ID: DND132_1480, Uniprot: F0JE79) (Figure 1), and has an ArsR-like HTH motif with a SAM-dependent methyltransferase similar to ArsM. Arsenite methyltransferase (ArsM) catalyzes the transfer of a methyl group to arsenite from S-adenosylmethionine (SAM) to form methylarsenicals, and is part of the ArsR regulon(69). Instead of methylarsenical production, ND132 ArsR4 is predicted to control methionine production and be responsive to SAH. No other homologues of ArsM are encoded in the genome of *P. mercurii* ND132. Like *arsR2* which is co-localized with *hgcAB*, *arsR4* is not part of an *ars* operon in *P. mercurii* ND132.

### Phylogenetic distribution of the ArsR2 *hgcAB* regulator

Close homologues of the ArsR2 regulator from *Pseudodesulfovibrio mercurii* ND132 are present in other *hgc*+ *Desulfovibrio* and *Pseudodesulfovibrio* genomes (Figure S1). To identify potential ArsR2 homologues, we searched for ArsR family proteins encoded before *hgcAB* in *Desulfovibrio* and *Pseudodesulfovibrio* reference genomes from a previous study(9). We also used the HMM of the HTH motif of ArsR (Pfam: PF01022, HTH_5.hmm) to broaden our search and identify all potential HTH family transcriptional regulators encoded in the reference genomes. We used protein sequence alignments and maximum likelihood phylogenies of HMM-hits to identify ArsR2 homologues. Among these genomes, we found ArsR2 homologues only in *hgc*+ *Desulfovibrionaceae* genomes, and always directly upstream of *hgcAB* (Figure S1). Other researchers working with a close relative to ND132, *Pseudodesulfovibrio hydrargi* BerOc1 demonstrated that these *arsR*-like genes and *hgcA* are co-expressed(8). Notably, all but one of the available *Pseudodesulfovibrio* genomes encode the ArsR2 homologue. However, co-occurrence of *arsR* with *hgcAB* does not appear to be phylogenetically conserved in the majority of *hgc*+ genomes of close relatives of *Desulfovibrionaceae* (Figure S1).

ArsR family regulators are also encoded with *hgcAB* in genomes from outside of the *Desulfovibrionaceae* family. In *hgc*+ genomes from the Hg-MATE-Db(7, 11), we identified putative ArsR/SmtB family regulators of *hgcAB* in isolate and environmental metagenome-assembled genomes (MAGs) from Deltaproteobacteria, Nitrospira, Spirochaetes, Ignavibacteriae(70) and Bacteroidetes(71), including *Paludibacter jiangxiensis* NM7(72) (Table S4). Based on protein sequence alignments, not all of these putative *hgcAB* regulators are close homologues of the ArsR2 protein from *Pseudodesulfovibrio mercurii* ND132. In some of these genomes, *hgcA* and *hgcB* are also colocalized with other *ars* genes (e.g. *arsC*, *arsB*, *arsM*). It is noteworthy that many of the *hgc*+ MAGs(73) of sediment and groundwater-associated bacteria from the Rifle IFRC site(74) have an ArsR-like regulator encoded with *hgcAB* and *ars* genes. This co-occurrence of *hgcAB* with *ars* genes in MAGs from arsenic-impacted environments has been noted previously(4, 43). For this study, we focused our analytical efforts on understanding possible ArsR-regulated transcription of *hgcAB* in *Pseudodesulfovibrio mercurii* ND132. However, more analysis is needed to explore a possible evolutionary relationship between the *ars* operon and Hg-methylation genes, and whether horizontal gene transfer of *hgcAB* may have been influenced by arsenic.

### Structural model

The ArsR2 homologues from *Pseudodesulfovibrio* and *Desulfovibrio* are phylogenetically distinct from other ArsR/SmtB family transcriptional regulators (Figure S2). The protein sequence alignment shows that ArsR2 has some of the conserved invariant residues associated with the HTH family of regulators (Figure S2)(75). To provide additional insight into the structure of the putative *hgcAB* transcriptional regulator (ArsR2, DND132_1054), we generated a homodimeric model of the protein using AlphaFold2(58, 59) (Figure 2). The cysteine-rich site associated with metal(loid) binding is different in ArsR2 compared to ArsR repressors of the *ars* operon. In *E. coli* ArsR (Uniprot: P15905), the As(III) binding site is associated with the three cysteines on the α3 helix (Cys32, Cys34, and Cys37)(64). The cysteines in ND132 ArsR2 (Uniprot: F0JBE8) occur on a coil after helix 2 and a beta strand before helix 4 (Figure S3). The ArsR2 from ND132 includes a cysteine (Cys26) near the N-terminus and four cysteines, Cys47, Cys49, Cys86, and Cys88 (Figure 2 and Figure S3) in each predicted metal(loid) binding site(75). Homologs from several other closely related bacteria of the *Pseudodesulfovibrio* and *Desulfovibrio* have similar structures and cysteine positions (Figure S4), suggesting that some of them may have similar functions. The four-cysteine motif in the metal(loid) binding site is conserved in all ArsR2 homologs found in *Pseudodesulfovibrio* and *Desulfovibrio* genomes (Figure S4) except for *Desulfovibrio brasiliensis* (WP_157054761.1), which has a three-cysteine motif.

**Figure 2.**
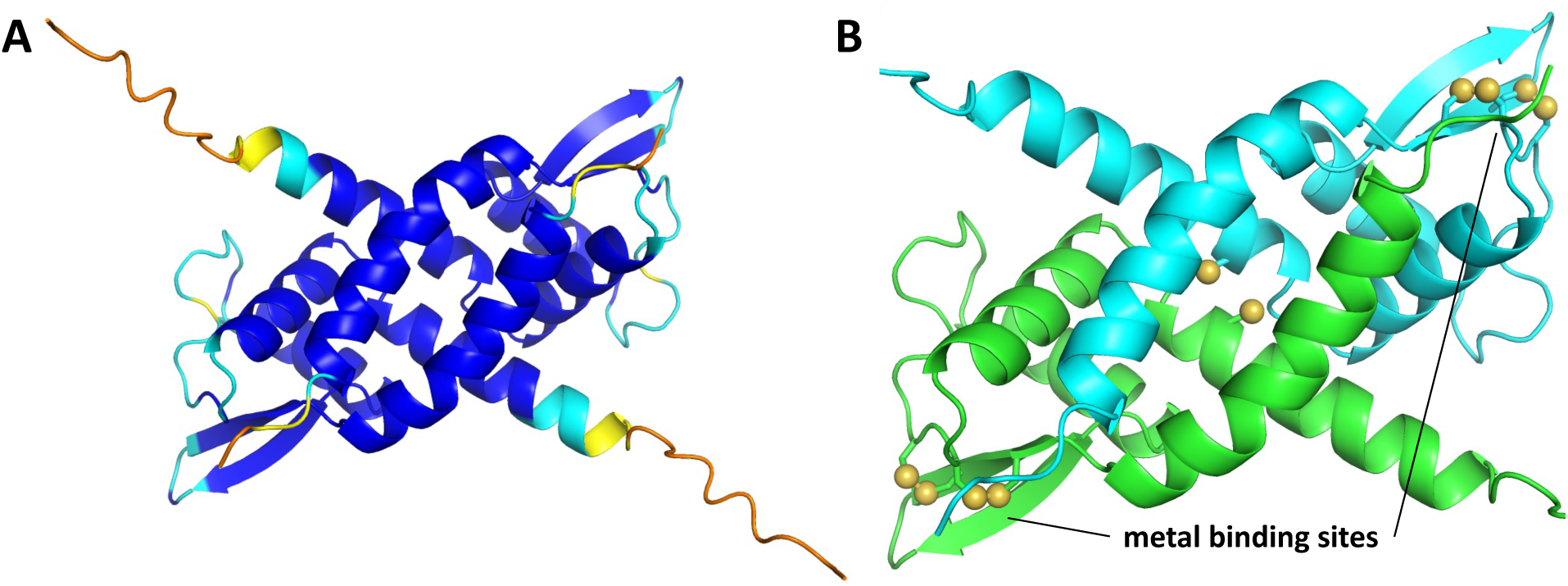
(A) AlphaFold2 model of the putative ArsR-like transcriptional regulator encoded by *arsR2* (Uniprot ID: F0JBE8) and co-localized with *hgcAB* in the genome of *Pseudodesulfovibrio mercurii* ND132. The model is colored by model confidence (i.e., predicted LDDT score). Dark blue = very high confidence (pLDDT > 90), cyan = confident (90 > pLDDT > 70), yellow = low confidence (70 > pLDDT > 50), and orange = very low confidence (pLDDT < 50). (B) Close-up view colored by chain with cysteine residues shown in the ball-and-stick representation.

The metal(loid) binding domain of the putative ArsR of *hgcAB* in *Pseudodesulfovibrio mercurii* ND132 (ArsR2, DND132_1054) differs from other ArsR repressors, containing four rather than three cysteines. Arsenite (As(OH)_3_) is predicted to coordinate with the three cysteine residues of ArsR as As(III), after the loss of its hydroxide ligands(75). However, only two of the three cysteines (Cys32, Cys34) are needed to bind arsenic to ArsR of *Escherichia coli* plasmid R773(76). The presence of four cysteine residues in the metal(loid) binding sites of the ArsR2 regulator suggests it may preferentially bind metal(loid) cations such as Zn(II), Cd(II), and Pb(II) like the four-cysteine residues in the binding motif of CadC(75). Alternatively, there could be 3-coordinate binding of As(III) to three of the four cysteine residues as has been proposed with As(III) and the SAM methyltransferase, ArsM(77).

### Arsenic tolerance in ND132

The presence of a putative ArsR regulator upstream of *hgcAB* suggests that the ability to methylate Hg may confer As resistance in some microbes. We compared As toxicity between wild-type (WT) ND132 and the *hgcAB* deletion mutant (ND132Δ*hgcAB*) to test the idea that the presence of *hgcAB* in the ND132 genome might contribute to As resistance. We performed growth assays in defined (no yeast extract) CCM-PF and CCM-LS media. Cultures were inoculated into media amended with sodium arsenate (0.2, 0.5, 1, 2, 5, or 20 mM) or sodium arsenite (0.02, 0.2, or 0.5 mM) and grown at 30 °C to stationary phase (Figures 3 and S5).

**Figure 3.**
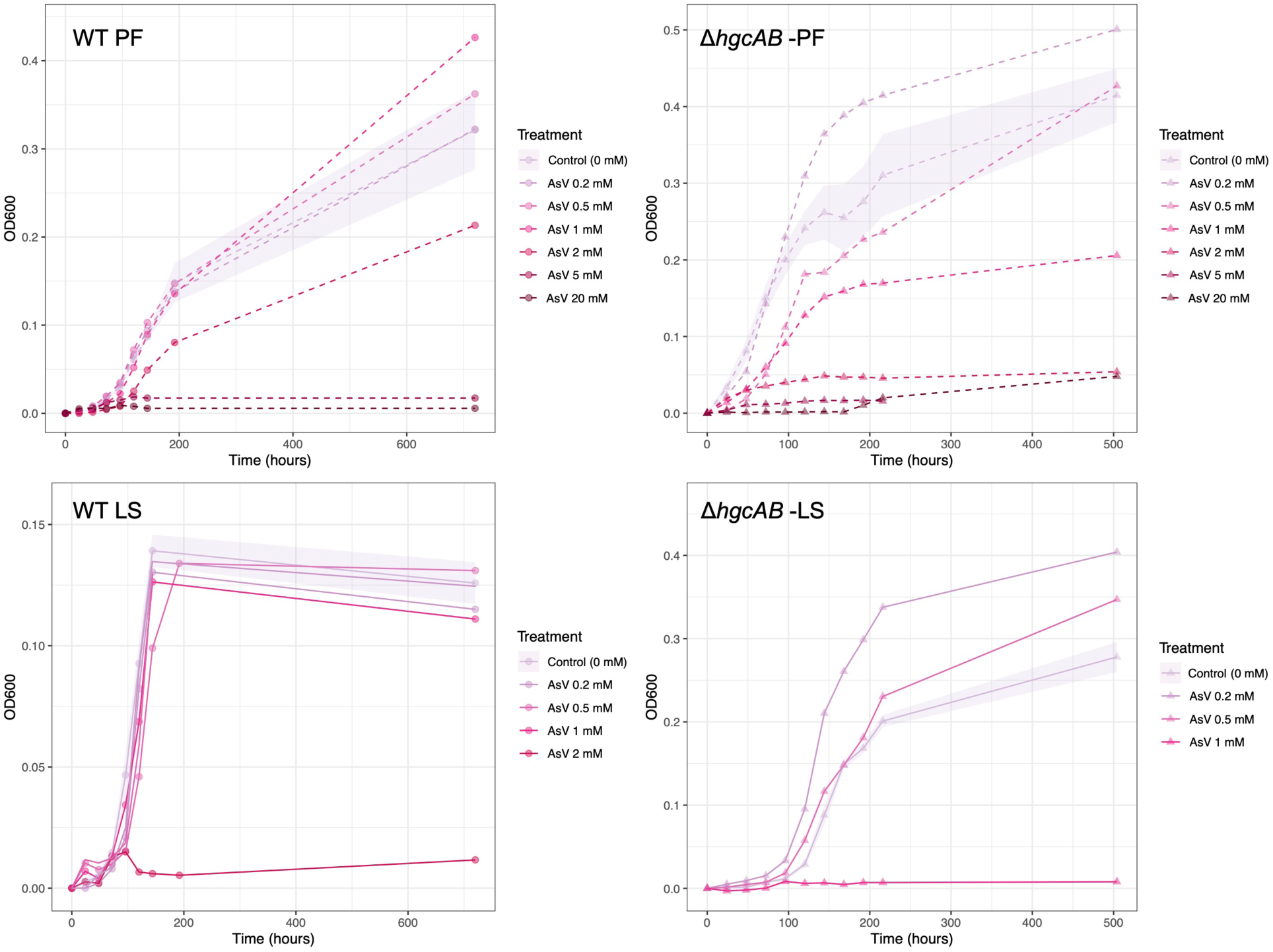
Growth assay tests of arsenate toxicity to *Pseudodesulfovibrio mercurii* ND132 wild-type (WT) and *hgcAB* deletion mutant (Δ*hgcAB*) in defined CCM media with pyruvate-fumarate (PF) or lactate-sulfate (LS) and sodium arsenate, As(V). Growth curves fitted to optical density measured at 600 nm (OD_600_) absorbance of biological triplicates using ‘ggplot’ and ‘dplyr’ in R. Shaded area shows the standard error of average OD_600_ measurements for control replicates.

The WT strain was more tolerant of As(V), than the knockout. Specifically, WT ND132 growth was unaffected by As(V) levels up to 1 mM on both media, while growth of ND132Δ*hgcAB* was significantly impacted at that level (Figure 3). Results were evaluated statistically by normalizing OD_600_ measurements to matched unamended controls, and then evaluating differences in normalized OD_600_ between the strains. Data from the exponential part of the growth curves were used in the statistical tests. Differences were assessed using general linear models that accounted for As concentration, time, and any significant interactions. The *hgcAB* knockout was significantly more sensitive to As(V) at levels above 0.2 mM on PF medium and at levels above 0.5 mM on LS medium (p <0.0001). Notably there was apparent hormesis in growth of ND132Δ*hgcAB*, with slightly enhanced growth at the lowest tested AsV levels.

The WT and knockout strains were equally resistant to As(III), although As(III) toxicity to both was roughly 5-10 times higher than As(V) (Figure S5). Culture chemistry, particularly the formation of As-sulfide complexes, impacted As toxicity. Both wild-type and ND132Δ*hgcAB* cultures had lower tolerance to both As species when grown under sulfidogenic conditions. A yellow mineral precipitate was observed when media with 0.5 mM As(III) was inoculated with sulfidogenic *P. mercurii* ND132 WT and Δ*hgcAB* cultures, and likely was the formation of orpiment (As_2_S_3_).

The reduced As(V) resistance of the *hgcAB* knockout strain suggests that *hgcAB* may confer some As(V) resistance. Or alternatively, the loss the *hgcAB* from the genome may impact other mechanisms of As(V) resistance by ND132. This observation deserves more exploration, especially the lack of differential resistance to the more toxic As(III).

### ND132 transformation of As(III) and As(V)

*Pseudodesulfovibrio mercurii* ND132 has several ArsR-regulated arsenic resistance operons (Figure 1). These operons are predicted to encode As(V) reductase, As(III) and methylarsenite permeases, and a putative organic arsenic resistance protein. To determine how ND132 may transform arsenic (As), we measured As speciation and mass balance in *P. mercurii* ND132 WT cultures grown with either 0.02 mM sodium arsenite or 0.2 mM sodium arsenate in defined CCM-PF media. At these concentrations, As did not affect cell growth rate (Figure S6A). We did not examine whether ND132 gains energy from As(V) reduction.

Under these conditions, we found that wild-type *P. mercurii* ND132 reduced As(V) to As(III) and produced potential thioarsenicals from both As(III) and As(V) (Figure S6, Table S5). As concentrations and speciation were measured initially and at the end of exponential growth (7 days). ND132 reduced about 25% of added As(V) to As(III) after 7 days, and converted about 3% to an unidentified As compound. This compound was tentatively identified as a thioarsenical based on the retention time of the peak (personal communication from Ben Wozniak at Brooks Applied). ND132 converted a higher fraction (roughly 15%) of added As(III) to the presumed thioarsenical compound (Figure S6).

In both As(III) and As(V) amended cultures, As speciation at the initial time point matched the spikes, and 70-80% of the added As could be accounted for after 7 days growth. Uninoculated medium served as abiotic controls. There was no significant loss of As or change in As speciation in abiotic controls, including no production of unidentified As compounds. No monomethyl- or dimethylarsenicals were detected in abiotic or cultured samples.

### Potential transcriptional regulation of *hgcA* by ArsR2

To help determine the transcriptional controls on the putative ArsR regulator of *hgcAB*, we challenged cells with As and SAH and evaluated expression of both *arsR2* and *hgcA*. We chose these potential regulators because, despite shared homology, the putative *hgcAB* regulator ArsR2 is distinct from other HTH regulators of the ArsR/SmtB family, and its function is unknown. Additionally, since the four other ArsR annotated in the genome of *P. mercurii* ND132 are predicted to regulate either methionine production or As resistance, we speculate that there could be biochemical overlaps between HgcAB function and As resistance or methionine production. We made this inference based on the co-localization of *hgcAB* with other *ars* genes(4, 43), and the shared homology of HgcA/B to corrinoid iron-sulfur proteins utilized in one-carbon metabolism(18, 78), including methionine biosynthesis. Thus, one goal of this work was to test the idea *hgcAB* may be part of the ArsR or SahR regulons. Because of the unusual configuration of the metal(loid) binding motif in ArsR2, we chose to test ND132 response to As(V) as well as As(III) and SAH.

We used real-time qPCR to test relative changes in expression of *arsR2* and *hgcA* during growth with potential transcription regulators (e.g., As(III), As(V), SAH). Tests were conducted in defined (i.e., no yeast) CCM media or yeast-enriched EPF media. Expression was normalized to three housekeeping genes and compared to control cultures without added treatments. We also tested for an effect of Hg on *hgcA* transcription and found none (Figure 4). This observation is consistent with previous work that *hgcA* expression and Hg-methylation activity are not induced by Hg(28–30). To evaluate a potential *arsR2* transcriptional response to SAH we tested several compounds in the methionine biosynthesis pathway. Because *P. mercurii* ND132 is not predicted to have an SAH uptake mechanism, we tested for transcriptional response to SAH by growing cells with methionine (1 mM), adenosine-2’,3’-dialdehyde (0.1 mM), and equimolar homocysteine with adenosine (0.2/0.2 mM, 0.5/0.5 mM, and 1/1 mM), which enabled the build-up of SAH in the cell. Adenosine-2’,3’-dialdehyde is an inhibitor of SAM-dependent methyltransferase. Methionine, homocysteine, and adenosine are metabolites in the SAM cycle, and higher concentrations in culture media should contribute to higher concentrations of SAH compared to cultures grown without these substrates. Growth of *P. mercurii* ND132 was completely inhibited at all concentrations of homocysteine with adenosine. Therefore, this treatment was not used for further expression or Hg methylation experiments.

**Figure 4.**
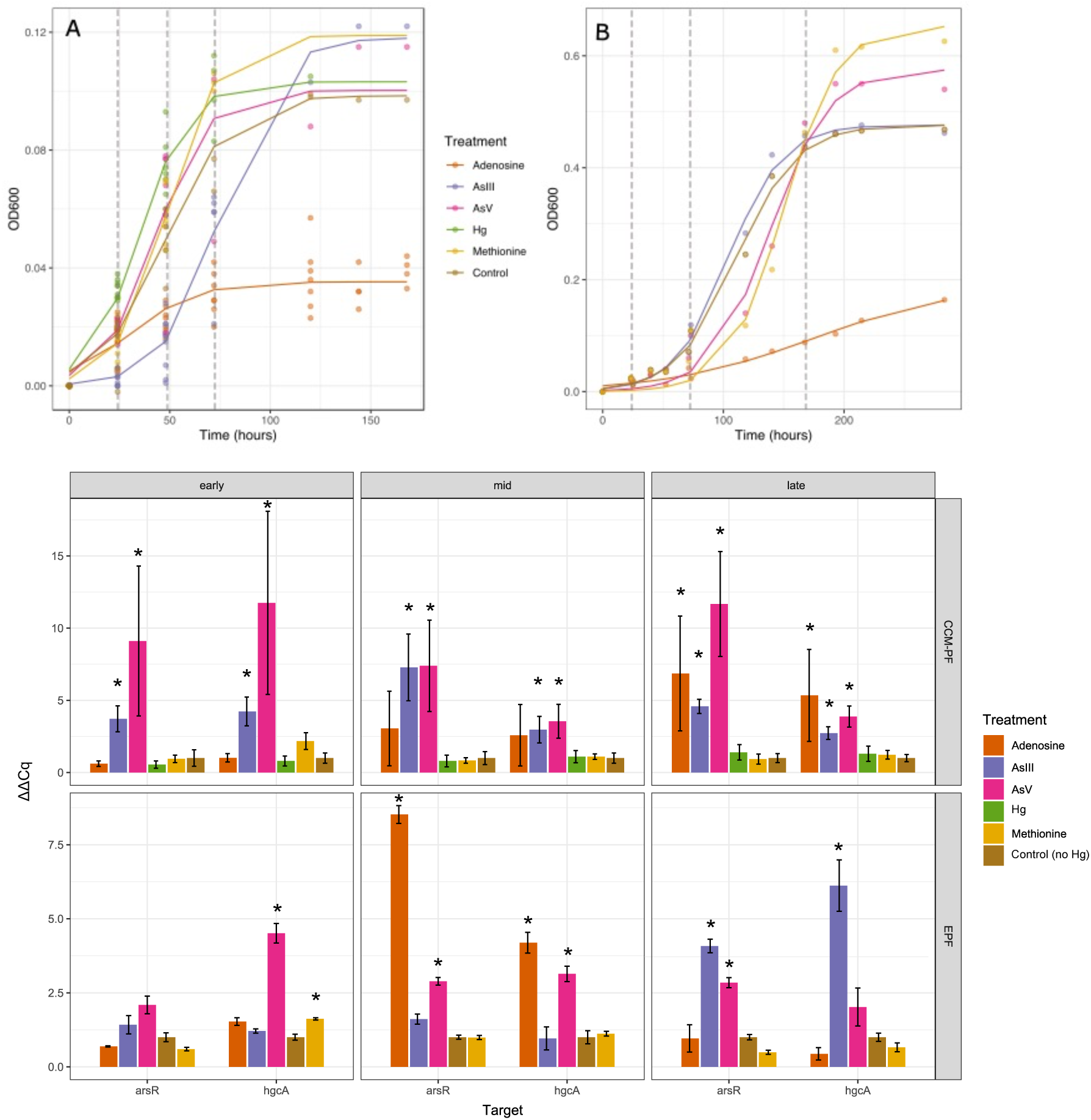
Regulation of *arsR* and *hgcA* expression in wild type *Pseudodesulfovibrio mercurii* ND132. Top: Growth curves of for cell grown in defined CCM-PF (A) and enriched EPF media (B) with various potential transcriptional regulators. Growth plotted as change in optical density at 600 nm absorbance (OD_600_) from inoculation at T_0_, using R program, Growthcurver(94). Gray dashed lines indicate when cells were harvested for RNA extraction for qPCR at early, mid, and late exponential growth. Bottom: Relative expression (ΔΔCq) of *arsR2* (DND132_1054) and *hgcA* (DND132_1056) from qPCR of cDNA (from RNA). Expression of the target genes was normalized to three housekeeping genes (*gyrA*, *gyrB*, *recA*) and to control (i.e., culture with no treatment). Treatments: 0.1 mM (CCM and EPF) adenosine-2’,3’-dialdehyde, 0.2 mM (CCM) and 0.02 mM sodium arsenite, 2.0 mM (CCM) and 0.2 mM (EPF) sodium arsenate, 1.0 mM HgCl (CCM only), 1.0 mM (CCM and EPF) methionine. Stars indicate significant difference between ΔΔCq of treatment compared to control (z-Test, p < 0.05). qPCR was performed on biological triplicate cultures (CCM) and single culture (EPF) and included three analytical replicates per cDNA sample.

### Transcription of *hgcA* is controlled by arsenic and SAH

We found that ArsR2 regulated *hgcA* expression in *P. mercurii* ND132 and can be responsive to both As and SAH. We observed significantly higher expression (z-test, one-tailed p-value < 0.05; Table S6) of *arsR2* and *hgcA* in *P. mercurii* ND132 cultures compared to control when grown with an SAH-hydrolase blocker (i.e., adenosine-2’,3’-dialdehyde) and with As. The transcriptional response varied depending on culture conditions (defined vs. enriched media), and growth phase (i.e., early, mid, or late exponential) (Figure 4). In defined CCM media, we observed significantly higher expression (Table S6) of *arsR2* and *hgcA* in cultures grown with As(V) and As(III) across the growth curve (Figure 3). We only observed significantly higher expression of *arsR2* and *hgcA* from adenosine dialdehyde at late exponential phase (Figure 4). Relative expression of *arsR2* and *hgcA* in enriched media differed from defined media conditions. In enriched media, we observed significantly higher *hgcA* expression in As(V)-amended cultures early in exponential growth, and later in exponential growth for As(III) amended cultures (Table S6). Compared to defined media, we observed significantly higher *hgcA* expression in methionine and adenosine amended cultures at earlier stages of growth (Figure 4). We attribute some of these differences to the presence of additional sulfur-containing metabolites in yeast extract (e.g., methionine) that could have contributed to increased SAH build-up in cultures in enriched media when grown with methionine or adenosine dialdehyde. Without yeast extract, the 1 mM methionine likely did not cause enough SAH build-up to induce a transcriptional response. Unlike in defined media conditions, *arsR2* transcriptional changes did not mirror *hgcA* expression patterns.

Differences in *hgcA* expression observed across the growth curve could indicate that the transcriptional response is regulated by an unknown substrate or by an As species produced by *P. mercurii* ND132. This effect could vary over the growth curve, as the substrate or products of As metabolism build up in culture or the cell over time. Arsenic speciation analyses showed that *P. mercurii* ND132 reduces As(V) to As(III), and produces an unknown thioarsenic species from As(III) (Figure S6). Differences in expression profiles between media could also be attributed to differences in As concentrations, with 10X less As(III) and As(V) used in enriched EPF media than defined CCM media. Reducing the amount of starting As(III) from 0.2 mM to 0.02 mM could have increased the amount of time needed to reach a threshold for transcriptional response. The lower As(III) concentration used in EPF media may explain why increased *arsR2* and *hgcA* expression was only observed in As(III)-amended EPF cultures at late exponential growth (Figure 4). The higher starting concentration of As(V) (0.2 mM) compared to As(III) (0.02 mM) in enriched EPF, may account for the earlier observed transcriptional response in As(V)-amended cultures. Overall, these results show that growing *P. mercurii* ND132 with As(III) or As(V) will significantly increase *hgcA* expression.

### Metabolites and media components can confound expression analysis

When cells were washed of culture media and then resuspended in basal salts with treatments, we observed significantly higher *arsR2* and *hgcA* expression due to As but not SAH (Figure 5). Washing cells of media components removed effects of adenosine dialdehyde on *arsR2* and *hgcA* expression observed in culture. Cells that were grown in enriched media with the various treatments were harvested at mid-log (OD_600_ = 0.2; Figure 4B), kept anaerobic, washed of media, and resuspended in a phosphate-salt buffer with the different treatments and 1 nM Hg. This was the same culture in which we measured *arsR2* and *hgcA* expression (Figure 4, EPF mid-log panel). The difference in observed *arsR2* and *hgcA* expression in washed cells (Figure 5C), compared to the culture (before washing) (Figure 4) indicates that build-up of SAH (or another metabolite) during growth with methionine and adenosine dialdehyde likely causes the observed transcriptional response of *arsR2* (DND132_1054). The transcriptional response to As, either as As(III) and or As(V), is more direct and instantaneous. We attribute differences in the transcriptional response to As over the growth curve as resulting from changes in As speciation and toxicity. The qPCR results from washed cells and batch cultures indicate that the *hgcA* regulator (DND132_1054) is responsive to both As and SAH.

**Figure 5.**
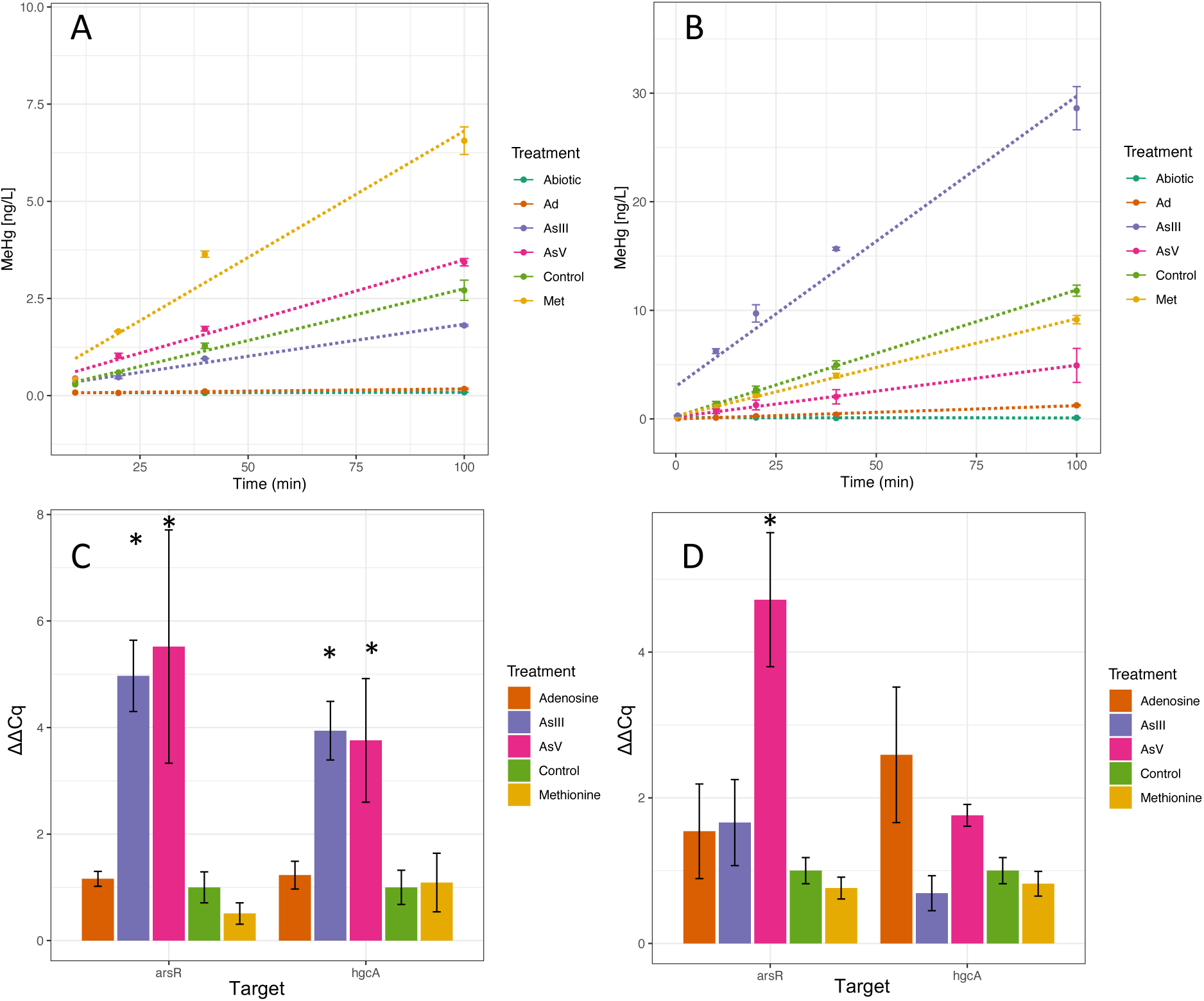
Impact of potential *hgcA* transcription regulators on MeHg production by *Pseudodesulfovibrio mercurii* ND132. Top: MeHg production in washed-cell (A) and batch cultures (B) with treatments: 0.1 mM adenosine-2’,3’-dialdehyde, 0.02 mM sodium arsenite, 0.2 mM sodium arsenate, 1.0 mM methionine, control (cells in unamended washed buffer or EPF media), and abiotic (no cells, unamended wash buffer or EPF media). Linear regression fit to data using R. Hg methylation assays were performed in triplicates, with sampling at 0, 10, 20, 40, 100 minutes after being spiked with 1 nM of ^201^HgCl. Error bars represent +/-1 standard error of the mean. Cells were harvested from assays at t = 100 minutes for qPCR of cDNA (from RNA) to measure normalized expression (ΔΔCq) of *arsR* (DND132_1054) and *hgcA* (DND132_1056) in washed cells (C) and batch culture (D). Expression normalized to three housekeeping genes (*gyrA, gyrB, recA*) and control (Supp. Methods 1.1). Stars indicate significant difference between ΔΔCq of treatment compared to control (z-Test, p < 0.05).

### As affects Hg methylation rates in ND132

We directly measured the impact of As and SAH on Hg methylation to help confirm that changes in *hgcAB* expression affected Hg methylation rates. Since *hgcA* transcription is regulated by *arsR2* in response to As and SAH, we expected As and SAH should also impact Hg methylation. Arsenic significantly affected *P. mercurii* ND132 Hg-methylation rates, but effects varied depending on whether cultures were washed of media components prior to methylation assays. We performed Hg-methylation assays in both cultures and washed cells using mid-log cultures (OD_600_ ∼ 0.2, Figure 4B) grown in EPF. Prior to methylation assays, cells were inoculated into media amended with observed expression regulators of *hgcA*: methionine, adenosine dialdehyde, As(III), or As(V). In washed cells harvested at mid-log phase, methionine and As(V)-treated cells produced significantly more MeHg than the control, while As(III) and adenosine significantly reduced MeHg production (Figure 5A). In unwashed cultures, Hg methylation rates were significantly higher in As(III)-amended cultures, but significantly lower in cultures grown with As(V) or adenosine, compared to control (Figure 5B). In batch culture assays, there was no significant difference in MeHg produced in cultures grown with methionine compared to control.

Differences in metabolic activity and biomass could account for some of the differences in MeHg production between treatments. When grown with As(V) and methionine, *P. mercurii* ND132 had a steeper growth curve and reached a higher optical density compared to control and other treatments (Figure 4B). In the washed-cell assays, the higher metabolic activity (CO_2_ respiration rate) (Figure S7) in the methionine treatment could explain some of the higher MeHg production (Figure 5A). In As(V)-treated washed cells, higher MeHg production could not be explained by differences in metabolic activity, biomass, or unfiltered Hg at T_0_ (Figure S8, Figure 6). Normalization of methylation rates to various measures of cell growth and metabolism did not change the relative impact of treatments (Figure 6). The measured initial concentration of total ^201^Hg was the same in all bottles, and Hg mass balance was maintained in all bottles (Figure S9). However, the filterable (0.22 μm filters) ^201^Hg concentration varied among the treatments. Nevertheless, normalization of methylation rates to initial filterable ^201^Hg concentrations did not alter the relative impact of treatments (Figure 6).

**Figure 6.**
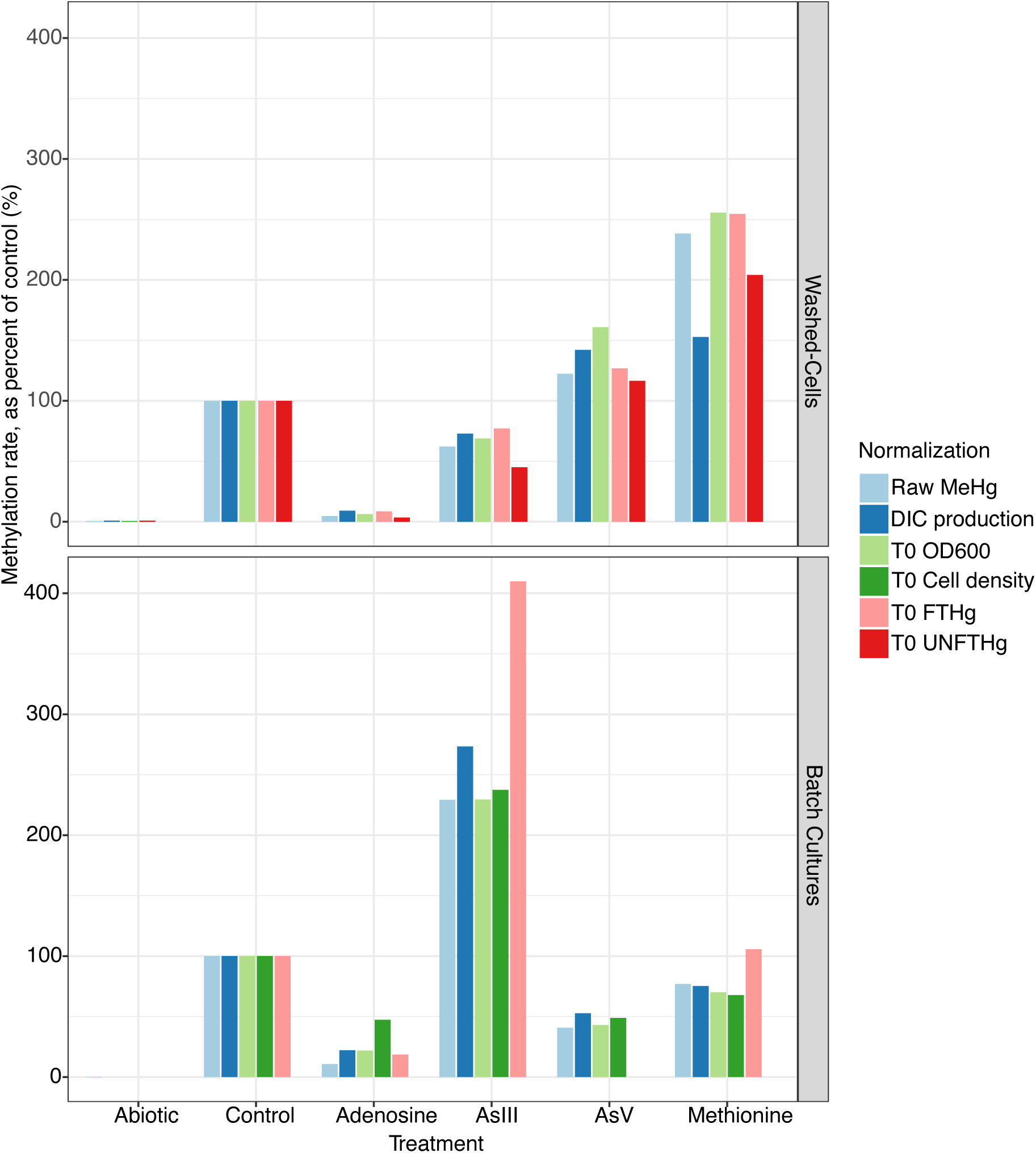
Normalized mercury methylation rates from culture and washed-cell assays. Mercury methylation rates in *Pseudodesulfovibrio mercurii* ND132 washed-cell (top panel) and batch culture (bottom panel) assays with test treatments compared to control when normalized to dissolved inorganic carbon (DIC) production (i.e., CO_2_ respiration), optical density (OD_600_) at T_0_, cell density at T_0_, filter-passable total Hg at T_0_, and total unfiltered Hg at T_0_.

In the Hg-methylation assays performed with mid-log cultures, significantly more MeHg was produced in the As(III) treatment (Figure 5B). This effect held true even when normalized to biomass (OD_600_, cell counts) (Figure S10), pH (Figure S11), metabolic activity (Figure S12), and total Hg (Figure S13) (Figure 5). Hg-methylation rates in As(V)-amended cultures remained significantly lower than control when normalized for these factors (Figure 6). In washed cells and culture media, adenosine-2’,3’-dialdehyde appears to limit cell growth and metabolic activity of *P. mercurii* ND132 (Figure 4). MeHg production in adenosine-amended washed cells and batch culture assays were only slightly higher than abiotic controls. It was difficult to decouple the effects of lower metabolic activity and less biomass on limited MeHg production. However, normalizing for biomass (as cell density) put Hg-methylation rates on par with As(V)-amended cultures in media (Figure 6).

Differences in Hg bioavailability may contribute to the differences in MeHg production among treatments. Sulfide has strong effects on Hg bioavailability for methylation(12, 30, 79, 80) and, although grown without sulfate, ND132 cleaves the S−C bond in cysteine provided in growth medium to release small amounts of sulfide, generally below 10 μM(51). We consistently measured significantly less sulfide in filtered samples from As(V)-amended washed-cell (Figure S7) and batch culture assays (Figure S11) compared to other treatments, including As(III). This observation indicates that As(V) impacts sulfide speciation, possibly by disrupting metabolic pathways that produce reduced sulfide, or precipitation of As species with sulfide. The higher concentrations of As(V) (0.2 mM) could account for the observed differences from As(III) (0.02 mM). In addition, the dissolved ^201^Hg spike is rapidly lost to cell walls and any precipitates in the medium. We observed differences in filterable ^201^Hg among treatments in both washed cells (Figure S9) and batch cultures (Figure S13). To test the impact of amendments on bioavailable Hg, we normalized methylation rates to the initial concentration of ^201^Hg in each treatment. Because Me^201^Hg is exported from cells after production(17), filterable total ^201^Hg concentrations after T_0_ often reflect Me^201^Hg production (Figure 6).

### Hg methylation rates decoupled from *hgcA* transcription

Increased *hgcAB* transcriptional activity did not explain increased MeHg production in most treatments. With the exception of the As(V)-washed cells, we did not observe significantly higher Hg methylation by cells with higher *hgcA* expression compared to control (Figure 5). Despite seeing significantly higher MeHg production by washed cells amended with methionine, and cultures grown with As(III), we did not observe significant differences in *hgcA* expression compared to control in qPCR analyses (Figure 5). Washed cells amended with As(III) had significantly higher *hgcA* expression (Figure 5), but produced significantly less MeHg (Figure 6). Similarly, cultures grown with adenosine-2’,3’-dialdehyde and As(V) treatments had higher expression of *hgcA*, but produced significantly less MeHg compared to control (Figure 5). This finding was surprising as we expected increased *hgcA* expression to result in higher Hg-methylation rates. However, *hgcA* activity may not have predicted higher Hg-methylation rates because of several factors that we could not account for in this experiment, 1) increased transcription of *hgcAB* may not have resulted in increased translation of HgcAB, 2) Hg availability and uptake by cells may be more rate-limiting than *hgcA*/HgcA activity, 3) Hg complexation with As may interfere with Hg availability for methylation(81), 4) possible competition between As and Hg for binding to HgcAB. Furthermore, the increased Hg methylation rate observed in As(III)-amended cultures could be from increased activity of other unknown proteins essential to the biochemical Hg methylation reaction.

### As and adenosine increase *hgcA* and *hgcB* expression in ND132 transcriptomes

RNA-seq analyses of RNA extracted from mid-log washed cells and batch cultures confirm that both *hgcA* and *hgcB* expression is regulated by As and SAH. Growing *P. mercurii* ND132 with As or adenosine dialdehyde appeared to have a significant influence on the transcriptome (Figure S14). We observed significantly higher expression (|log FC| > 1.5, p < 0.05) of *arsR2* (DND132_1054), *hgcA* (DND132_1056) and *hgcB* (DND132_1057) in As(III) and As(V)-treated washed cells compared to control (Figure 7). These features were also significantly differentially expressed (p < 0.05) in batch cultures amended with As(V) and adenosine dialdehyde (Figure S15). The RNA-seq data show that *hgcB* is being up-regulated with *hgcA*, which is important since both genes are needed for Hg methylation(19). Overall, there were more significant shifts in batch culture transcriptomes compared to washed-cells, with many of the same features being differentially expressed across treatments (Figure S14). Methionine amended batch cultures had the least significant shift in its transcriptome compared to control (n = 4; Figure S16). In *P. mercurii* ND132 cultures grown with adenosine, the predicted SahR regulator encoded by *arsR4* (Gene ID: DND132_1481) was significantly upregulated (log FC = 2.36, p = 2.64 E-19) along with the adenosylhomocysteinase-encoding gene *ahcY* (Gene ID: DND132_1480, log FC = 2.31, p = 2.16E-16) (Figure S16). The SAH responsive regulator was not significantly upregulated in any other transcriptome, including methionine-treated cultures. This observation indicates that there may not have been enough build-up of SAH in *P. mercurii* ND132 cultures amended with 1 mM methionine to induce a transcriptional response from SahR or ArsR. There appears to be a cross-reactivity to SAH and As by two ArsR encoded in the genome of *P. mercurii* ND132 *arsR2* (DND132_1054) as well as *arsR5* (DND132_2311) (Figure S15). However, exposure to As(III) or As(V) did not induce a transcriptional response by SahR (*arsR4*, DND132_1481).

**Figure 7.**
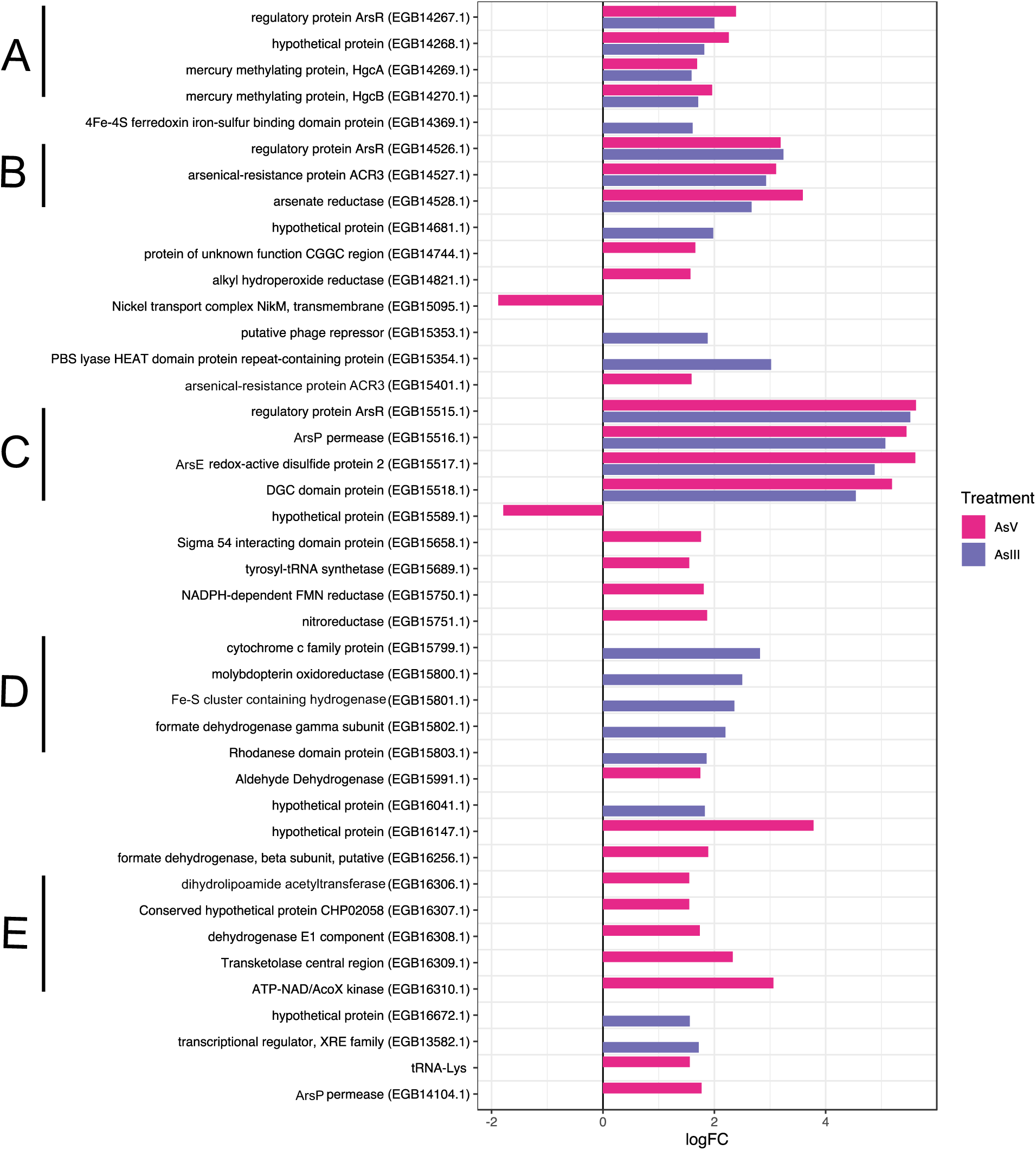
Differential expression of significant features (|log FC| > 1.5, p < 0.05) from RNA-seq analyses of As(V) (0.2 mM) and As(III) (0.02 mM) treated washed cells of wild-type *Pseudodesulfovibrio mercurii* ND132. Several putative operons (A-E) had significant differential expression compared to control washed-cells (i.e., no arsenic treatment).

### Co-expression of *hgcAB* with operons of arsenic resistance

*Pseudodesulfovibrio mercurii* ND132 is resistant to As(V) and As(III), and may also be resistant to organoarsenicals and respire As(V). Both Hg-methylation genes (*hgcA* and *hgcB*) appear to be part of the ArsR regulon and are co-regulated with *ars* operons. Based on RNA sequencing, there were 12 shared features (Figure S14) that had significant differential expression (|log FC| > 1.5, p-value < 0.05) in As-treated washed cells compared to control (Figure 7). These features were also significantly differentially expressed (p < 0.05) in batch cultures amended with As(III), As(V), and adenosine dialdehyde (Figure S15). In addition to increased *hgcA* and *hgcB* expression, two ArsR-regulated *<u>ars</u>*operons had significantly higher expression in As-amended washed cells and batch cultures (Figure 7, Figure S15). Operon B (Figure 8) encodes for ArsR3, an Acr3 As(III) efflux permease, and arsenate reductase. Another set of upregulated genes (Operon C) encodes ArsR5 and enzymes homologous to those utilized in organoarsenical resistance. Operon C was also upregulated in the transcriptome of adenosine dialdehyde cultures. The other ArsP (DND132_0889) and Acr3 (DND132_2196) permeases encoded outside an operon were not consistently upregulated with the *ars* operons (Figure 7, Figure S15). The only *arsR* gene of *P. mercurii* ND132 that was not significantly differentially expressed in any transcriptome was *arsR1* (DND132_0890).

**Figure 8.**
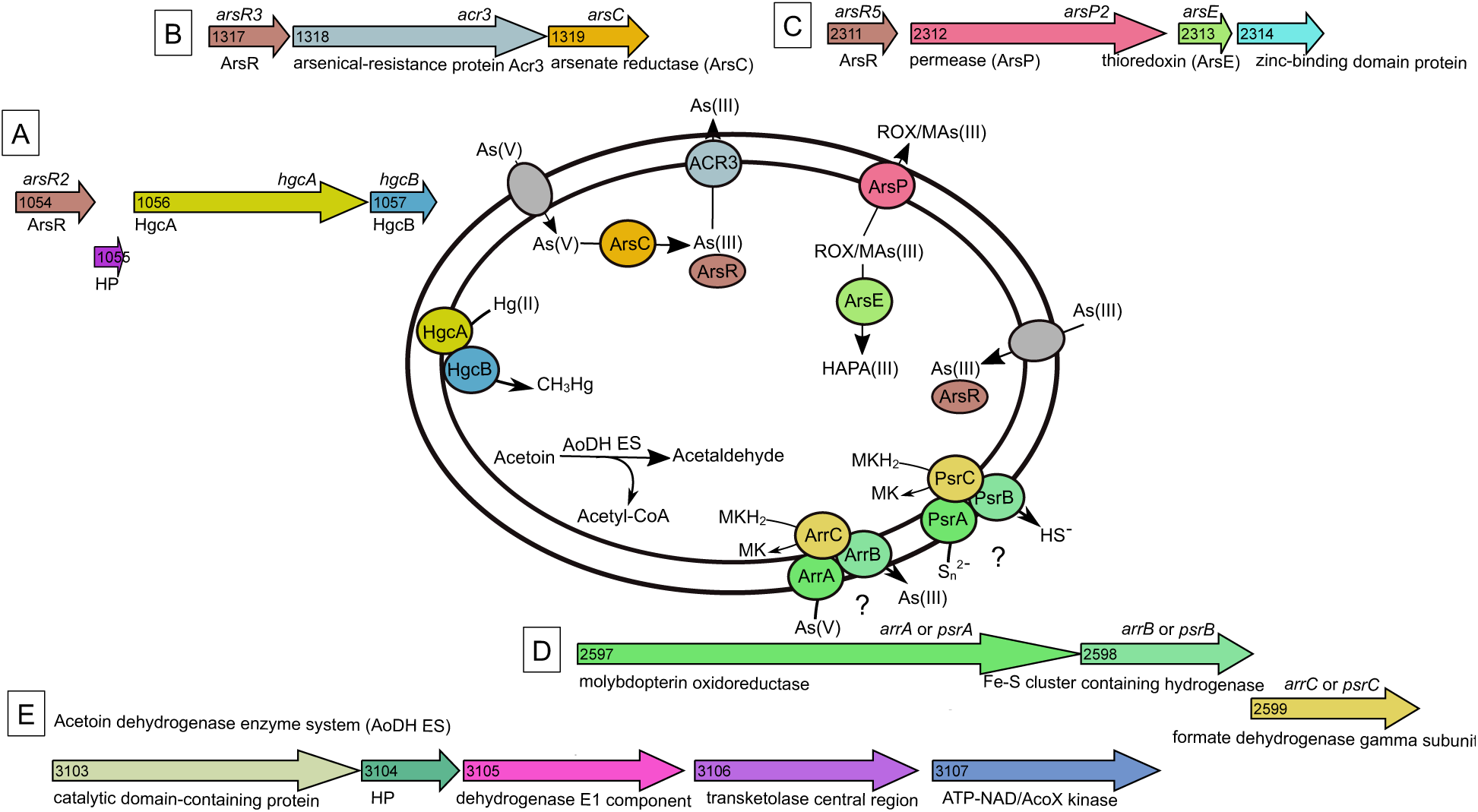
Proposed biochemical reactions for operons that were significantly differentially expressed in *Pseudodesulfovibrio mercurii* ND132 transcriptomes grown with arsenic. This includes genes that are part of the ArsR regulon in ND132, including (A) Hg-methylation genes *hgcA* and *hgcB,* (B) operon that encodes for arsenate reductase (ArsC) and As(III) efflux pump Acr3, and (C) an operon that includes homologues of organoarsenic resistance proteins ArsE and ArsP. Also significantly differentially expressed was an operon (D) that encodes homologues of the polysulfide reductase (Psr) or arsenic respiration complex (Arr), which couples menaquinone (MK) oxidation to periplasmic AsV or sulfur reduction, and (E) a set of genes that encode homologues of the acetoin dehydrogenase enzyme system (AoDH ES). *P. mercurii* ND132 is a confirmed Hg-methylator and can reduce As(V) to As(III). The biochemical mechanism that produces the unknown thioarsenic compound is unknown. To date, *P. mercurii* ND132 has not been tested for organoarsenic resistance or for acetoin degradation. The mechanism for uptake of As(V) and As(III) by ND132 is unknown. Operon graphic made using Gene Graphics(93).

In As(III) washed cells and As(V) and adenosine batch cultures, an operon with possible links to arsenic or polysulfide respiration was upregulated compared to control. The set of genes (Operon D, Figure 8) encodes possible homologues of the As(V) respiratory reductase (Arr) or polysulfide reductase (Psr) complexes. Since transcription of these genes was increased by As(III), we speculated that their encoded function may relate to As respiration. The genes encode homologues of ArrA (DND132_2597), ArrB (DND132_2598), and ArrC (DND132_2599), and are respectively annotated in *P. mercurii* ND132 as a molybdopterin oxidase, an Fe-S cluster-containing hydrogenase, and formate dehydrogenase gamma subunit protein. The Arr complex is related to the DMSO-molybdopterin (DMSO-MPT) reductase family, with Arr subunit A having a Mo/W-*bis*(pyranopterin guanosine dinucleotide) cofactor in its active site and a [4Fe-4S] center, and Arr subunit B with four [4Fe-4S] clusters(82). A diverse range of microorganisms can respire As(V) for energy using ArrAB(83), including *Chrysiogenes arsenatis*, *Shewanella* strain ANA-3, *Alkaliphilus oremlandii* and *Geobacter* sp. Strain OR-1(84, 85). In some microorganisms, ArrAB can function as a bidirectional enzyme(86), oxidizing As(III) to As(V). Often, ArrAB are encoded in microbial genomes alongside genes of the *ars* operon. However, in *P. mercurii* ND132 the putative operons are not co-localized. The ND132 ArrA homolog includes the conserved molybdenum and Fe-S binding domains found in known functional respiratory As(V) reductases but is missing some of the key active-site residues(86).

Instead, operon D (Figure 8) may encode for polysulfide or thiosulfate reduction rather than As(V) respiration. Based on the phylogenetic relationship between the putative ArrA in *P. mercurii* ND132 with other DMSO-MPT family proteins, the protein encoded by DND132_2597 is more closely related to bacterial polysulfide reductase (PsrA) than ArrA from known As(V)-respiring bacteria (Figure S17). Microbes in deep-sea vents and hydrothermal springs use the membrane protein complex PsrABC to couple periplasmic polysulfide or thiosulfate reduction with quinone oxidation(87). The operon in *P. mercurii* ND132 also encodes homologues of cytochrome *c* oxidase (DND132_2596) and two rhodanese-domain proteins (DND132_2600, DND132_2601), one of which (DND132_2600) is significantly upregulated in As(III) cells. A second cluster of putative *psrAB* genes (DND132_1314, DND132_1313) are present elsewhere in the genome (Figure S17), and likely encode for cytoplasmic sulfur reduction(88). The transcription of these *psrAB* genes was not significantly affected by As or SAH. The DMSO-MPT family of proteins that includes Psr and Arr has been implicated in thioarsenic production. The geothermal isolate *Pyrobaculum yellowstonensis* strain WP30 produces thioarsenate during respiratory elemental sulfur and As(V) reduction(89). More work is needed to know if the protein complex encoded by operon D interacts with As(III), As(V), or with thiosulfates/polysulfides and enables *P. mercurii* ND132 to respire As. However, it may be responsible for producing the unknown thioarsenic species observed in *P. mercurii* ND132 culture media.

In As(V) washed cells an operon with homology to the acetoin dehydrogenase complex was upregulated. In batch cultures, this operon was significantly downregulated in all treatments except methionine (Figure S15). The operon is predicted to encode an acetoin dehydrogenase enzyme system (AoDH ES) that catalyzes the conversion of acetoin to acetyl-CoA and acetaldehyde (Figure 8). The AoDH ES contains multiple copies of three enzymatic components: acetoin oxidoreductase (E1), dihydrolipoamide acetyltransferase (E2) and dihydrolipoyl dehydrogenase (E3)(90, 91). The operon (Operon E, Figure 8) encodes AoDH ES homologues annotated as a dihydrolipoamide acetyltransferase (DND132_3103), (PFam PF00198, PF00364, PF02817), an acetoin:2,6-dichlorophenolindophenol oxidoreductase subunit alpha (DND132_3105) (dehydrogenase E1 component, PFam PF00676), acetoin:2,6-dichlorophenolindophenol oxidoreductase subunit beta (DND132_3106) (transketolase central region, PFam PF02779, PF02780), and an acetoin catabolism protein with ATP-NAD/AcoX kinase motifs (PFam PF01513). These proteins also share homology to enzymes in the pyruvate dehydrogenase complex that convert pyruvate into acetyl-CoA by pyruvate decarboxylation(92). The operon is possibly regulated by a sigma(54)-dependent HTH transcriptional regulator (DND132_3102). In closely related *Pseudodesulfovibrio hydrargi* BerOc1 this protein is annotated as an acetoin dehydrogenase operon transcriptional activator AcoR (Uniprot: A0A1J5NC98). No obvious link to As metabolism or resistance could be found. It is possible that this putative AoDH ES operon is upregulated in *P. mercurii* ND132 As(V) washed cells because of cross-reactivity of As with the HTH regulator (DND132_3102) or because of a knock-on effect from other differentially regulated mechanisms.

### Summary

The broad research aim of these experiments was to examine transcriptional regulation of *hgcAB* in *Pseudodesulfovibrio mercurii* ND132, with a focus on the potential role of an upstream ArsR-like regulator found in *hgc*+ Deltaproteobacteria. Additionally, we examined the relationship between Hg methylation rates in *Pseudodesulfovibrio mercurii* ND132 with metabolic activity and *hgcAB* transcriptional activity. We have shown that *hgcAB* expression in *Pseudodesulfovibrio mercurii* ND132 is transcriptionally regulated by an ArsR-like regulator encoded by *arsR2* (DND132_1054) that is responsive to As and SAH. Since *P. mercurii* ND132 can reduce As(V) to As(III), and we observed increased *hgcA* and *hgcB* expression when grown with both species, it is likely that As(III) coordinates with ArsR2. The *hgcAB* transcriptional regulator shares homology to other members of the ArsR/SmtB family, but has a distinct four-cysteine motif in its metal(loid) binding site. This homology also distinguishes it from other ArsR family regulators encoded in the genome of *P. mercurii* ND132. Since As(III) preferentially binds to three cysteine residues in ArsR repressors of the *ars* operon, we cannot exclude the possibility that another As species or heavy metal(loid) cation may coordinate with ArsR2 and also control *hgcAB* expression. Although the transcriptional response to As was more pronounced, under some culture conditions methionine and adenosine dialdehyde also significantly increased *arsR2* and *hgcAB* transcription. We attribute this transcriptional response to a build-up in SAH within the cell.

The RNA-seq analyses show that *hgcAB* transcription is co-regulated as part of the ArsR regulon with operons that encode As resistance. We also observed significant co-expression of *hgcAB* with genes that encode putative As(V) or polysulfide respiration when *P. mercurii* ND132 is grown with As. These pathways may explain the possible thioarsenic production observed in *P. mercurii* ND132 cultures. This indicates that As biogeochemistry may influence *hgcAB* expression and Hg-methylator activity in some environmental settings. In this study, we did not observe a significant loss in As resistance with a loss in *hgcAB* function in *Pseudodesulfovibrio mercurii* ND132. However, the co-expression and co-localization of *hgcAB* with arsenic resistance genes in some *hgc*+ genomes may hold clues to *hgcAB* evolution and possible environmental drivers of horizontal transfer of these genes. More work is needed to tease apart possible overlaps between HgcAB function and arsenic resistance and respiration mechanisms.

Results from these experiments do not confirm that higher *hgcAB* expression alone increases MeHg production, but that growth and culture conditions, and the nature of *hgcAB* transcriptional regulation probably all play roles in methylation control. Both As(III) and As(V) impacted Hg methylation and transcription of *hgcAB* and *arsR* in *P. mercurii* ND132, although the impacts were not always coupled. We also observed that metabolic activity and microbial biomass can significantly influence Hg methylation rates in cultures, and counter possible impacts of *hgcAB* activity. Hg methylation rates may also be impacted by HgcAB translation, protein folding, co-factor and membrane insertion, and the methyl donor mechanism. Combining information of *hgcAB* transcription with microbiome composition, metabolic activity, and Hg speciation and geochemistry may help resolve environmental Hg methylation rates. It is therefore important to have accurate and reliable methods for measuring these factors and using multiple parameters when incorporating microbial dynamics into predictive models for Hg methylation.

## ACKNOWELDGEMENTS

Caitlin Gionfriddo was a Robert and Arlene Kogod Secretarial Scholar with the Smithsonian Environmental Research Center while conducting work described in this manuscript. This work was funded by the U.S. Department of Energy, Office of Science, Office of Biological and Environmental Research, Subsurface Biogeochemical Research (SBR) Program, and is a product of the Critical Interfaces Science Focus Area at Oak Ridge National Laboratory, which is managed by UT-Battelle, LLC under Contract No. DE-AC05-00OR22725 with the U.S. Department of Energy. The As speciation analysis was performed by Brooks Applied Labs and we thank Ben Wozniak and Collette Machado for technical support. The RNA-seq transcriptome analyses were performed at Azenta US Inc. and we thank Andrew Corbin and Eric Danzeisen for technical support.

## DATA AVAILABILITY STATEMENT

The transcriptomic sequencing data is publicly available through KBase narrative (https://narrative.kbase.us/narrative/124642).

## SUPPLEMENTARY MATERIAL

See attached document of supplementary methods, tables, and figures.

