## Supplemental Materials for "Transcriptional control of *hgcAB* by an ArsR*-*like regulator in *Pseudodesulfovibrio mercurii* ND132"

**Supplementary Materials for "Transcriptional control of *hgcAB* by an ArsR-like regulator in *Pseudodesulfovibrio mercurii* ND132"**

Caitlin M. Gionfriddo*[1], Ally Bullock Soren[1], Ann Wymore[2], D. Sean Hartnett[1], Mircea Podar[2], Jerry M. Parks[2], Dwayne A. Elias[3], Cynthia C. Gilmour[1]

### Supplemental Methods

### 1. Mercury Analytical Methods

***1. 1. Total Hg analysis.*** Isotope-specific total Hg concentrations in filtered and unfiltered cultures and media were measured using a modification of EPA Method 1631 (EPA-821-R-01-023, Mar 2001), using dual CVAF and ICP-MS (Agilent 7900 ICP-MS) detection. , A Tekran (Toronto, Canada) 2600 purge and trap system was used for sample introduction to the ICP-MS following $SnCl_2$ (J. T. Baker) reduction to $Hg^0$. Samples were collected in PETG bottles and preserved at collection in 0.5% trace-metal grade HCl (Macron Fine Chemicals). For analysis, samples were placed in 40 ml clear VOA vials (Thomas Scientific), brought up to ~25 ml (if needed) with ultra-pure water (made in house using a re-circulating system that includes charcoal filtration, deionization, R/O and UV oxidation and brings resistance to >22 MΩ), and digested overnight at room temperature with 0.5% v/v concentrated BrCl per vial. Immediately prior to analysis, 0.1 ml 30% hydroxylamine hydrochloride (Acros Organics) was added to remove free chlorine. Samples were reduced with 0.1 ml 20% $SnCl_2$ (J.T. Baker). Reagents were prepared according to Method 1631, however, for BrCl, KBr and $KBrO_3$ (both (Beantown Chemicals, Hudson, NH) powders were muffled at $300^0$C to reduce Hg contamination. The hydroxylamine hydrochloride was purged with $N_2$ for 30 min after the addition of $SnCl_2$ to 0.1%, to reduce Hg content, and stored refrigerated. Blank subtraction was based on analysis of the sample matrix. Primary Hg standards were obtained from Brooks Rand at 1 ppm in 2% $HNO_3$. Working 1 ppb dilutions were made in 0.5% BrCl and stored in the refrigerator. Two additional NIST-traceable quality control (QCS) standards, from SCP Science (Montreal, Quebec, Canada) and Inorganic Ventures (Christianburg, VA), were used in each run. A dilution of NIST 1641e was used as a CRM.

***1.2 MeHg analysis.*** Isotope-specific MeHg concentrations were determined using a modification of EPA Method 1630 (EPA-821-R-01-020, Jan 2001) with dual ICP-MS and CVAF detection. Samples were cleaned up prior to analysis by steam distillation(1). A Tekran (Toronto, Canada) 2700 purge and trap GC system was used for ethylation and sample introduction to the ICP-MS. Samples were distilled in 60 ml fluoropolymer plastic (PFA) distillation jars (Savillex) in an aluminum heating block at 120 °C. Prior to distillation, samples were brought up to 22.5 ml (if needed) with ultra-pure water and amended with 0.2 ml 20% KCl (source), 2 ml of 0.2 M $CuSO_4$ (VWR) and 1 ml of 9M $H_2SO_4$ (Millipore). Distillates were trapped in a wet ice bath. The target distillation percentage was 65-70%. Flow rate was 45 ml/min $N_2$. Sample and distillate weights were recorded. The distillation gear was cleaned by scrubbing the stills with 409® cleaner and then boiling with 20% HCl.

For analysis, distillate aliquots were brought up to 27 ml with ultrapure water in 40 ml amber VOA vials (Thomas Scientific) and then buffered by the addition of 0.03 ml 5 mM Na acetate. -MeHg was derivatized by aqueous-phase ethylation with the addition of 0.03 ml 1% sodium tetraethylborate (Strem). Reagents were generally prepared according to Method 1630.

Ongoing Performance and Recovery (OPR) and QCS MeHg standards (1ppm in 0.5 % (v/v) acetic acid, 0.2 % (v/v) HCl) were obtained from Brooks Rand Instruments (Seattle, WA) yearly and held refrigerated. Two separate Brooks Rand standards were used as OPR and QCS standards. Working 1 ppb stock solutions were kept refrigerated in 0.5% trace-metal grade HCl, 0.5% acetate buffer and 10 μM glutathione to limit degradation. MeHg standards were tested against certified inorganic Hg standards before and after digestion to inorganic Hg with BrCl, to monitor any degradation over time(2). NIST 1566b oyster tissue(3) was used as the certified reference material (CRM). For MeHg samples with an experimental isotope spike, CRM recovery was used to correct MeHg recovery in samples.

***1.3 Synthesis of MeHg standards***. $Me^{201}Hg$ standards were synthesized in-house using methylcobalamin (Hintelmann and Ogrinc 2003; Bancon-Montigny 2004). Isotopically-enriched $^{201}Hg$ was purchased from ORNL in 1995 (the isotopic abundance was <0.02% $^{196}Hg$, 0.08% $^{198}Hg$, 0.1% $^{199}Hg$, 0.45% $^{200}Hg$, 98.11% $^{201}Hg$, 1.18% $^{202}Hg$, and 0.08% $^{204}Hg$. $CH_3{}^{201}HgCl$ was synthesized by dissolving 5 mg of methylcobalamin (Sigma-Aldrich) in 5 ml 4 M acetate buffer (pH 5) and adding 2 ml of a roughly 250 μg/ml stock solution of $^{201}Hg$ preserved in 0.5% HCl. The solution was held in the dark at 37 $^0$C for 2 h, then cooled immediately to 4 $^0$C. The synthesized $Me^{201}Hg$ was cleaned up by extraction with methylene chloride. Specifically, the solution was acidified with 1 ml of concentrated HCl, brought up to 10 ml with DI water, then extracted by shaking with 10 ml $MeCl_2$ for 5 min in a Teflon separation funnel. This step was repeated 5 times to produce a total of ~50 ml $MeCl_2$ extract. The extract was rinsed twice with 10 ml of ultra-pure water in another separation funnel. Finally, the $Me^{201}HgCl$ was brought back into the aqueous phase by heating the $MeCl_2$ with 10 ml of ultra-pure water to just below the boiling point of $MeCl_2$ (~40 $^o$C) under a stream of $N_2$ gas. Once the $MeCl_2$ was visibly gone, the residual water was heated for an additional 10 minutes to ensure that all of the $MeCl_2$ was removed. The final solution was acidified with 0.1 ml of 50% HCl and stored frozen in aliquots. The final concentration of $Me^{201}Hg$ was verified by isotope dilution ICP-MS.

***1.4 Hg and MeHg QA/QC.*** Results are reported as concentrations of excess $^{201}THg$ and $Me^{201}Hg$ after correction for background THg or MeHg. Detection limits were based on 3 times the standard deviation of the sample matrix blanks, adjusted to sample volume. We routinely analyzed roughly 10% duplicate samples (where volume allowed), ongoing and recovery standards, and a variety of blanks as part of our quality assurance and quality control (QA/QC) efforts (Table SI-1).

### 2. Quantitative PCR methods

#### *2.1 Calculations for normalized expression (ΔΔCq):*

List of terms and abbreviations:

E = efficiency

Cq = quantitation cycle (i.e. cycle in which fluorescence can be detected), mean of all technical and biological replicates, also called mean cycle of expression

Ctrl = Control

GOI=gene of interest

Sd = standard deviation

Equation 1. Relative Quantification (RQ)

$RQ_{GOI} = E^{(CqCtrl-CqGOI)}$

$RQ\ sd = sdCq_{GOI} * RQ_{GOI} * LN€$

Equation 2. Normalization Factor (NF)

$NF = (RQ_{gyrA} * RQ_{gyrB} * RQ_{recA})$^(1/3)

$NF\ sd = NF*SQRT((RQsd_{gyrA}/RQ_{gyrA}*3)^2+(RQsd_{gyrB}/RQ_{gyrB}*3)^2+(RQsd_{recA}/RQ_{recA}*3)^2)$

Equation 3. Normalized Expression (ΔΔCq)

$\Delta\Delta Cq_{GOI} = RQ_{GOI}/NF$

$\Delta\Delta Cq\ s.d. = \Delta\Delta Cq_{GOI}*(SQRT((s.d.RQ_{GOI}/RQ_{GOI})^2+(s.d.NF_{HKG}/NF_{HKG})^2)$

**3. Hg methylation assays in washed cells**

***3.1 Approach:*** *Pseudodesulfovibrio mercurii* ND132 was grown in EPF medium containing the various treatments, along with no amendment controls with and without cells. Cells were grown to mid-log phase, 40 mL of cells were aliquoted into triplicate conical tubes, pelleted at 2,800 x g for 15 min. Pellets were rinsed with wash buffer to remove extracellular metabolites. Cells were resuspended in 50 mL of minimal medium wash buffer containing 10 mM KCl, 171 mM NaCl,

4.4 mM $KH_2PO4$, 7.5 mM $NH_4Cl$, 5 mM MOPS buffer, 1 mM Na-fumarate, 1 mM Na-pyruvate, pH adjusted to 7.5 with NaOH, and reduced with 0.5 mM cysteine-HCl. Resazurin dye was used in wash buffer as a redox indicator. Wash buffer also contained the following test treatments: 0.1 mM adenosine 2'3'-dialdehyde, 1 mM methionine, 0.2 mM sodium arsenate, or 0.02 mM sodium arsenite. Control condition had no additional amendments. Methylation assays were carried out over 100 min after a 1 nM $^{201}$HgCl spike was added to resupended cells. We followed filterable and unfiltered Hg concentrations, pH, OD, sulfide concentration and $CO_2$ production through the assays.

***3.2 Treatment effects on metabolic activity and culture chemistry.*** There were initial differences in cell density of the washed cell cultures at $T_0$ among treatments (Fig S7), reflecting differences in culture growth, with Control=Met>AsIII=AsV>Ad, based on Student's least square means difference test with $\alpha$=0.050. Cultures were active in all of the treatments during the methylation assays, based on $CO_2$ production (Fig. S6), although OD did not increase significantly during the assays (Fig. S7).

Adenosine-2'3'-dialdehyde significantly reduced the growth and metabolic activity of ND132 based on several measures, including inorganic C production, optical density and sulfide production. Other treatments affected growth and metabolism but to a much lesser extent. In general, methionine (Met) cultures grew slightly better than controls, while AsV and AsIII cultures grew slightly worse.

Measured pH was consistent across treatments, and did not change during the methylation assays

(Fig. S7). Sulfide ranged from about 2 to 10 μM initially, and did not decline significantly during the 100 min assays.

***3.3 Hg methylation results.*** In this washed cell assay, Met significantly enhanced MeHg production relative to all other treatments, while AsIII and Ad slightly depressed methylation relative to unamended controls (Fig. 4). Normalization of methylation rates to various measures of cell growth and metabolism did not change the relative impact of treatments (Fig 5). In this study, the 1 nM $^{201}$Hg was rapidly lost to cells (Fig. S8). Only a small fraction of the 1 nM spike was recovered in the filterable fraction immediately after the spike. There was also some variability in the measured initial concentration of total unfiltered $^{201}$Hg (Fig S8). However, there was no loss of total $^{201}$Hg during the assays. Nevertheless, normalization of methylation rates to initial filterable $^{201}$Hg concentrations or to total $^{201}$Hg concentrations did not alter the relative impact of treatments (Fig 5).

**4. Hg methylation assays in EPF cultures**

***4.1 Approach.*** Assays were carried out using mid-log phase cultures grown on EPF medium with the various treatments. The culture was aliquoted into triplicate, 50 mL conical tubes for methylation assays. Methylation assays were carried out over 100 min after the addition of a 1 nM $^{201}$Hg spike. We followed filterable and unfiltered Hg concentrations, pH, OD, cell counts, sulfide concentration, short -chain fatty acids and $CO_2$ production.

***4.2 Treatment effects on metabolic activity and culture chemistry.*** There were initial differences in cell density at T0 among treatments (Fig S9), with Met>AsIII=Control>AsV>Ad, based on a

Tukey least square means test with $\alpha=0.050$. Cultures were active in all of the treatments during the methylation assays, based on $CO_2$ production (Fig. S11) and significant increases in OD and cell counts (Fig. S9). Adenosine-2'3'-dialdehyde significantly reduced the growth and metabolic activity of *P. mercurii* ND132 based on several measures, including inorganic C production, optical density, cell counts, and fumarate metabolism relative to control. Other treatments affected growth and metabolism but to a much lesser extent. In general, methionine and AsV cultures grew slightly better than controls, and AsIII grew slightly worse. *P. mercurii* ND132 cultures utilize fumarate and pyruvate to produce succinate and acetate. There were significant differences in the concentrations of these organic acids among the treatments at the beginning of the Hg methylation assays, with Adenosine-2'3'-dialdehyde treated cultures utilizing less fumarate and producing less succinate and acetate. However, changes in the concentrations of these organic acids were not measurable over the 100 min incubations. Medium chemistry varied slightly among treatments (Fig. S10): pH ranged from 6.6 to 6.8, sulfide ranged from 0.5 to 8 μM initially, declining to <4 μM in all treatments after 100 min. As we often find, sulfide levels were significantly lower in the AsV treatments than in controls.

**Supplemental Table S1**. QA/QC for Hg and MeHg analyses. Detection limits were based on 3X the standard deviation of process blanks adjusted for sample volumes. UNFMeHg = unfiltered MeHg. FTHg = filtered total Hg. Volumes in the washed cell study were too low for analytical replicates. The CRM for total Hg was a dilution of NIST 1641e. The CRM for MeHg was NIST 1566b oyster tissue.

| **Analysis** | **Experiment** | **Detection limit ± Stdev** | **Analytical replicates RPD ± Stdev** | **QCS standard recovery ± Stdev (%)** | **CRM recovery Avg ± Stdev (%)** |
|---|---|---|---|---|---|
| FTHg | Washed Cells | 0.02± 0.008 | | 102.2 ± 4.0 | 87.3 ± 7.1 |
| FTHg | Batch Culture | 0.055± 0.02 | 4.7 ± 1.7% | 99.5 ± 2.7 | 79.1 ± 1.3 |
| UNFMeHg | Washed Cells | 0.008 ± 0.004 | | 95.2 ± 7.8 | 61 ± 2.2 |
| UNFMeHg | Batch Culture | 0.043± 0.018 | 5.6 ± 4.8% | 100.9 ± 6 % | 55 ± 2.5 |

**Supplemental Table S2**. Custom designed primers for this study used in quantitative PCR of *arsR2* (before *hgcA*), *hgcA*, and three house-keeping genes, gyrase subunit A (*gyrA*) and B (*gyrB*), recombinase A (*recA*) in *Pseudodesulfovibrio mercurii* ND132

| **Primer Name** | **Target Size (nt bp)** | **Gene locus/ID** | **Sequence 5' to 3'** |
|---|---|---|---|
| DND132_1054-arsR-F | 180 | DND132_1054 | CGA CGA CGA GAC GTT CCT TG |
| DND132_1054-arsR-R | 180 | DND132_1054 | ACC AGA CCG GCA TTC TTG AG |
| DND132_1056-hgcA-F | 107 | DND132_1056 | GCC AAC TAC AAG CTG ACC TTC |
| DND132_1056-hgcA-R | 107 | DND132_1056 | CCC GCC GCG CAC CAG ACG TT |
| DND132_1603-recA-F | 153 | DND132_1603 | AAG TCC AAC TGC GTG GTC AT |
| DND132_1603-recA-R | 153 | DND132_1603 | GTC CTT GAG GGT CTG GAT GC |
| DND132_2285-gyrA-F | 148 | DND132_2285 | GCA CCA TCG AAT CGC TGA TG |
| DND132_2285-gyrA-R | 148 | DND132_2285 | CTC CTC CAG CTC GAC CAC GC |
| DND132_2284-gyrB-F | 182 | DND132_2284 | CCA AGA AGC TCA TCC AGA AG |
| DND132_2284-gyrB-R | 182 | DND132_2284 | CCT CGA AGA AGG TGT TCA GC |

**Supplemental Table S3**. Pairwise sequence identity (%) of *Pseudodesulfovibrio mercurii* ND132 ArsR proteins. Protein sequence alignment performed using MUSCLE(4) in Geneious Prime (v. 2022.0.1). ArsR proteins numbered by order of appearance in genome, with gene name (e.g. DND132_0890) also provided for reference.

| | **ArsR1 DND132_0890** | **ArsR2 DND132_1054** | **ArsR3 DND132_1317** | **ArsR4 DND132_1481** | **ArsR5 DND132_2311** |
|---|---|---|---|---|---|
| **ArsR1 DND132_0890** | | 28.947 | 22.901 | 26.606 | 23.894 |
| **ArsR2 DND132_1054** | 28.947 | | 23.308 | 25.714 | 32.727 |
| **ArsR3 DND132_1317** | 22.901 | 23.308 | | 22.936 | 26.050 |
| **ArsR4 DND132_1481** | 26.606 | 25.714 | 22.936 | | 18.947 |
| **ArsR5 DND132_2311** | 23.894 | 32.727 | 26.050 | 18.947 | |

**Supplemental Table S4**. List of potential Hg-methylators from the Hg-MATE-Db(5, 6) that have an ArsR/SmtB family regulator encoded in proximity of *hgcAB*. ArsR/SmtB family regulators were identified using HMM model (Pfam: PF01022, HTH_5.hmm) and cross-referencing HMM-hits with *hgcAB* loci.

| Organism | Genome ID (NCBI or JGI Gold) |
|---|---|
| Bacterium BMS3Abin13 | BDTF01000099.1 |
| Bacterium BMS3Bbin14 | BDTX01000009.1 |
| Bacteroidales bacterium 45-6 | MKRS01000096 |
| Bacteroidales bacterium isolate fen_945 | PMNI01000131 |
| Bacteroidales bacterium isolate fen_956 | PMIL01000100 |
| Bacteroidales bacterium isolate fen_957 | PMIJ01000040 |
| Bacteroidales bacterium isolate fen_959 | PMEL01000085 |
| Bacteroidales bacterium isolate fen_963 | PMBO01000363 |
| Bacteroidales bacterium isolate fen_974 | PLNT01000008 |
| Bacteroidales bacterium isolate fen_975 | PLNJ01000261 |
| Bacteroidales bacterium isolate fen_976 | PLNF01000075 |
| Bacteroidales bacterium isolate fen_977 | PLMT01000013 |
| Bacteroidales bacterium isolate fen_978 | PLMP01000132 |
| Bacteroidales bacterium isolate fen_980 | PLML01000094 |
| Bacteroidales bacterium isolate fen_982 | PLMC01000050 |
| Bacteroidales bacterium isolate fen_983 | PLLX01000153 |
| Bacteroidales bacterium isolate fen_984 | PLLO01000018 |
| Bacteroidales bacterium UBA1800 | DCFQ01000041 |
| Bacteroidales bacterium UBA4133 | DFWN01000090 |
| Bacteroidetes bacterium GWA2_30_7 | MENC01000050 |
| Bacteroidetes bacterium GWA2_31_9 | MEND01000285 |
| Bacteroidetes bacterium GWA2_32_17 | MENF01000219 |
| Bacteroidetes bacterium GWF2_35_48 | MEOG01000010 |
| Bacteroidetes bacterium HGW-Bacteroidetes-21 | PHDH01000120 |
| Bacteroidetes bacterium isolate PowLak16_MAG39 | QYPB01000007 |
| Bacteroidetes bacterium isolate SZUA-422 | QKCK01000329 |
| Bacteroidetes bacterium RIFOXYA12_FULL_35_11 | MEPE01000019 |
| Bacteroidetes bacterium RIFOXYC12_FULL_35_7 | MEPL01000424 |
| Balneolia bacterium isolate CSSed10_409R1 | PWPG01000342 |
| Deltaproteobacteria bacterium CG11_big_fil_rev_8_21_14_0_20_49_13 | PCWZ01000018 |
| Deltaproteobacteria bacterium HGW-Deltaproteobacteria-18 | PHBE01000009.1 |
| Deltaproteobacteria bacterium isolate B144_G9 | QMMX01000194.1 |
| Deltaproteobacteria bacterium isolate B3_G2 | QMMT01000133.1 |
| Deltaproteobacteria bacterium RIFOXYD12_FULL_57_12 | MGTE01000083 |

| | |
|---|---|
| Deltaproteobacteria bacterium SG8_13 | LJNK01000005.1 |
| Deltaproteobacterium sp. OalgD3 | Ga0117946 |
| Deltaproteobacterium sp. OalgD4 | Ga0117947 |
| Desulfobacteraceae bacterium isolate B1Sed10_16 | PUOB01000242.1 |
| Desulfobacteraceae bacterium isolate Glo_11 | PIVS01000212.1 |
| Desulfobacteraceae bacterium isolate Glo_9 | PIVQ01001248.1 |
| Desulfobacteraceae bacterium isolate maxbin2.1429 | RPPU01000027.1 |
| Desulfobacteraceae bacterium isolate metabat2.783 | RPRI01000233.1 |
| Desulfobacteraceae bacterium isolate T3Sed10_121 | PWYD01000231 |
| Desulfobacteraceae bacterium UBA5616 | DIKO01000105.1 |
| Desulfobacterales bacterium RIFOXYA12_FULL_46_15 | MGTJ01000099.1 |
| Desulfobacterales bacterium SG8_35_2 | LJTL01000115.1 |
| Desulfobulbaceae bacterium isolate BM004 | PKTW01000107.1 |
| Desulfobulbaceae bacterium isolate BM004 sc_3649 | PKTW01000107 |
| Desulfobulbaceae bacterium isolate SZUA-575 | QKIH01000002.1 |
| Desulfobulbaceae bacterium UBA2248 | DDXY01000062.1 |
| Desulfobulbaceae bacterium UBA4056 | DFZM01000029.1 |
| Desulfobulbaceae bacterium UBA5750 | DIFK01000123 |
| Desulfobulbaceae bacterium UBA5750 UBA5750_contig_8051 | DIFK01000123.1 |
| Desulfobulbus sp. Tol-SR | JROS01000055.1 |
| Desulfocurvus vexinensis DSM 17965 | JAEX01000001.1 |
| Desulfohalovibrio alkalitolerans DSM 16529 | ATHI01000029.1 |
| Desulfoluna spongiiphila strain AA1 | FMUX01000014.1 |
| Desulfomicrobium apsheronum strain DSM 5918 | FORX01000017 |
| Desulfomicrobium escambiense DSM 10707 | AUAR01000028.1 |
| Desulfomicrobium norvegicum strain DSM 1741 | FOTO01000006 |
| Desulfomicrobium sp. UBA5193 | DHWB01000004.1 |
| Desulfonatronum lacustre DSM 10312 | KI912608.1 |
| Desulfonatronum sp. SC1 | PZKN01000012.1 |
| Desulfonatronum thioautotrophicum strain ASO4-1 | KN882169.1 |
| Desulfonatronum thiodismutans strain MLF-1 | JPIK01000009.1 |
| Desulfonatronum thiosulfatophilum strain ASO4-2 | FMXO01000006.1 |
| Desulfonatronum zhilinae Al915-01 | Ga0139010 |
| Desulfosarcina cetonica JCM 12296 | BBCC01000140.1 |
| Desulfospira joergensenii DSM 10085 | ATUG01000002.1 |
| Desulfotignum balticum DSM 7044 | ATWO01000001.1 |
| Desulfotignum phosphitoxidans DSM 13687 | APJX01000008.1 |
| Desulfovibrio ferrophilus | AP017378.1 |
| Desulfovibrio profundus strain 500-1 | LT907975.1 |

| | |
|---|---|
| Desulfovibrio sp. isolate SP109 | PARK01000001.1 |
| Desulfovibrio sp. UBA1335 | DBST01000087.1 |
| Desulfovibrio sp. UBA6079 | DIXN01000049 |
| Desulfovibrio sp. UBA6079 | DIXN01000049.1 |
| Desulfovibrio sp. UBA7315 | DKQF01000002.1 |
| Desulfovibrio sp. X2 | ATHV01000019.1 |
| Desulfovibrionaceae bacterium UBA6814 | DKEU01000210.1 |
| Firmicutes bacterium HGW-Firmicutes-9 | PGZM01000005 |
| Ignavibacteriae bacterium isolate fen_1254 | PMML01000079 |
| Ignavibacteriae bacterium isolate fen_1261 | PMNH01000227 |
| Ignavibacteriae bacterium isolate fen_1272 | PMKT01000182 |
| Ignavibacteriae bacterium isolate fen_1278 | PMJP01000036 |
| Ignavibacteriae bacterium isolate fen_1287 | PMDB01000064 |
| Ignavibacteriae bacterium isolate fen_1292 | PMBK01000005 |
| Ignavibacteriae bacterium isolate fen_1298 | PLNH01000092 |
| Ignavibacteriae bacterium isolate fen_1299 | PLMS01000101 |
| Ignavibacteriae bacterium isolate fen_1300 | PLMK01000008 |
| Nitrospira bacterium SG8_35_1 | LJTK01000093.1 |
| Paludibacter jiangxiensis NM7 | BDCR01000003 |
| Pseudobacteroides cellulosolvens DSM 2933 | LGTC01000001 |
| Pseudodesulfovibrio aespoeensis Aspo-2 | CP002431.1 |
| Pseudodesulfovibrio hydrargyri strain BerOc1 | LKAQ01000001 |
| Pseudodesulfovibrio indicus strain DSM 101483 | SOBK01000001.1 |
| Pseudodesulfovibrio indicus strain J2 | CP014206.1 |
| Pseudodesulfovibrio mercurii ND132 | CP003220.1 |
| Spirochaetae bacterium HGW-Spirochaetae-1 | PGXY01000003 |
| Spirochaetae bacterium HGW-Spirochaetae-5 | PGXT01000070 |
| Spirochaetes bacterium RBG_16_49_21 | MIBJ01000045 |
| Spirochaetia bacterium UBA5550 | DINC01000107 |
| Spirochaetia bacterium UBA5763 | DIEX01000034 |
| Unclassified Desulfobacterales bin111 | Ga0224686 |
| Unclassified Desulfovibrionales bin158 | Ga0224688 |

**Supplemental Table S5**. Arsenic speciation data from *P. mercurii* ND132 wild-type cultures (ND132-WT) grown in CCM media amended with 0.02 mM sodium arsenite or 0.2 mM sodium arsenate. Arsenic speciation measured in culture media before inoculation (T0) and at late exponential growth (T168). Abiotic controls are uninoculated media. Standard deviation (s.d) of the mean of two biological replicates. 'Unknown' are sum of all unidentified arsenic species.

| Culture | Arsenic Amendment | Sample Time (hr) | Species | mM | s.d. |
|---|---|---|---|---|---|
| ND132-WT | AsIII | 0 | AsIII | 0.0191 | 1.89E-04 |
| | | | AsV | 2.56E-04 | 7.55E-06 |
| | | | Unknown | b.d.l* | b.d.l* |
| ND132-WT | AsIII | 168 | AsIII | 0.0111 | 9.25E-04 |
| | | | AsV | 1.06E-04 | 2.07E-05 |
| | | | Unknown | 2.92E-03 | 2.17E-04 |
| Abiotic | AsIII | 0 | AsIII | 0.0196 | 0.00E+00 |
| | | | AsV | 2.92E-04 | 2.74E-05 |
| | | | Unknown | b.d.l* | b.d.l* |
| Abiotic | AsIII | 168 | AsIII | 0.0196 | 1.89E-04 |
| | | | AsV | 4.08E-04 | 1.60E-05 |
| | | | Unknown | b.d.l* | b.d.l* |
| ND132-WT | AsV | 0 | AsIII | b.d.l† | b.d.l† |
| | | | AsV | 0.215 | 0.0151 |
| | | | Unknown | b.d.l† | b.d.l† |
| ND132-WT | AsV | 168 | AsIII | 0.045 | 0.0033 |
| | | | AsV | 0.128 | 9.44E-05 |
| | | | Unknown | 4.74E-03 | 3.87E-04 |
| Abiotic | AsV | 0 | AsIII | b.d.l‡ | b.d.l‡ |
| | | | AsV | 0.198 | 0.0123 |
| | | | Unknown | b.d.l† | b.d.l† |
| Abiotic | AsV | 168 | AsIII | b.d.l‡ | b.d.l‡ |
| | | | AsV | 0.204 | 3.78E-03 |
| | | | Unknown | b.d.l† | b.d.l† |

*Analytical detection limit 6.7E-05 mM
†Analytical detection limit 3.3E-03 mM
‡Analytical detection limit 3.3E-03 mM

**Supplemental Table S6.** Normalized expression (ΔΔCq) of *arsR* (DND132_1054) and *hgcA* (DND132_1056) from qPCR of *P. mercurii* ND132 cultures grown with test treatments. Only conditions with significant difference in expression compared to control are included, based on one-tail p-value from z-test of means with variance of ΔΔCq.

| Media | Growth Stage | Target | Treatment | ΔΔCq | s.d. | z-score | p-value |
|---|---|---|---|---|---|---|---|
| CCM-PF | early | arsR | AsIII | 3.72 | 0.9 | 2.24 | 1.2E-02 |
| | | | AsV | 9.11 | 5.19 | 3.38 | 3.6E-04 |
| | | hgcA | AsIII | 4.23 | 1 | 2.89 | 1.9E-03 |
| | | | AsV | 11.75 | 6.34 | 4.16 | 1.6E-05 |
| CCM-PF | mid | arsR | AsIII | 7.28 | 2.31 | 4.12 | 1.9E-05 |
| | | | AsV | 7.39 | 3.16 | 3.62 | 1.5E-04 |
| | | hgcA | AsIII | 2.97 | 0.92 | 1.75 | 4.0E-02 |
| | | | AsV | 3.55 | 1.17 | 2.07 | 1.9E-02 |
| CCM-PF | late | arsR | Adenosine | 6.86 | 3.98 | 3.02 | 1.3E-03 |
| | | | AsIII | 4.58 | 0.49 | 4.80 | 7.9E-07 |
| | | | AsV | 11.67 | 3.63 | 5.63 | 9.2E-09 |
| | | hgcA | Adenosine | 5.34 | 3.18 | 2.34 | 9.6E-03 |
| | | | AsIII | 2.73 | 0.44 | 2.08 | 1.9E-02 |
| | | | AsV | 3.88 | 0.73 | 2.91 | 1.8E-03 |
| EPF | early | hgcA | AsV | 4.51 | 0.33 | 5.35 | 4.3E-08 |
| | | | Methionine | 1.62 | 0.04 | 1.66 | 4.9E-02 |
| EPF | mid | arsR | Adenosine | 8.52 | 0.3 | 12.36 | 0.0E+00 |
| | | | AsV | 2.89 | 0.13 | 4.23 | 1.2E-05 |
| | | hgcA | Adenosine | 4.19 | 0.35 | 4.23 | 1.2E-05 |
| | | | AsV | 3.14 | 0.26 | 3.09 | 1.0E-03 |
| EPF | late | arsR | AsIII | 4.08 | 0.23 | 5.44 | 2.6E-08 |
| | | | AsV | 2.84 | 0.17 | 3.61 | 1.5E-04 |
| | | hgcA | AsIII | 6.12 | 0.87 | 5.09 | 1.7E-07 |

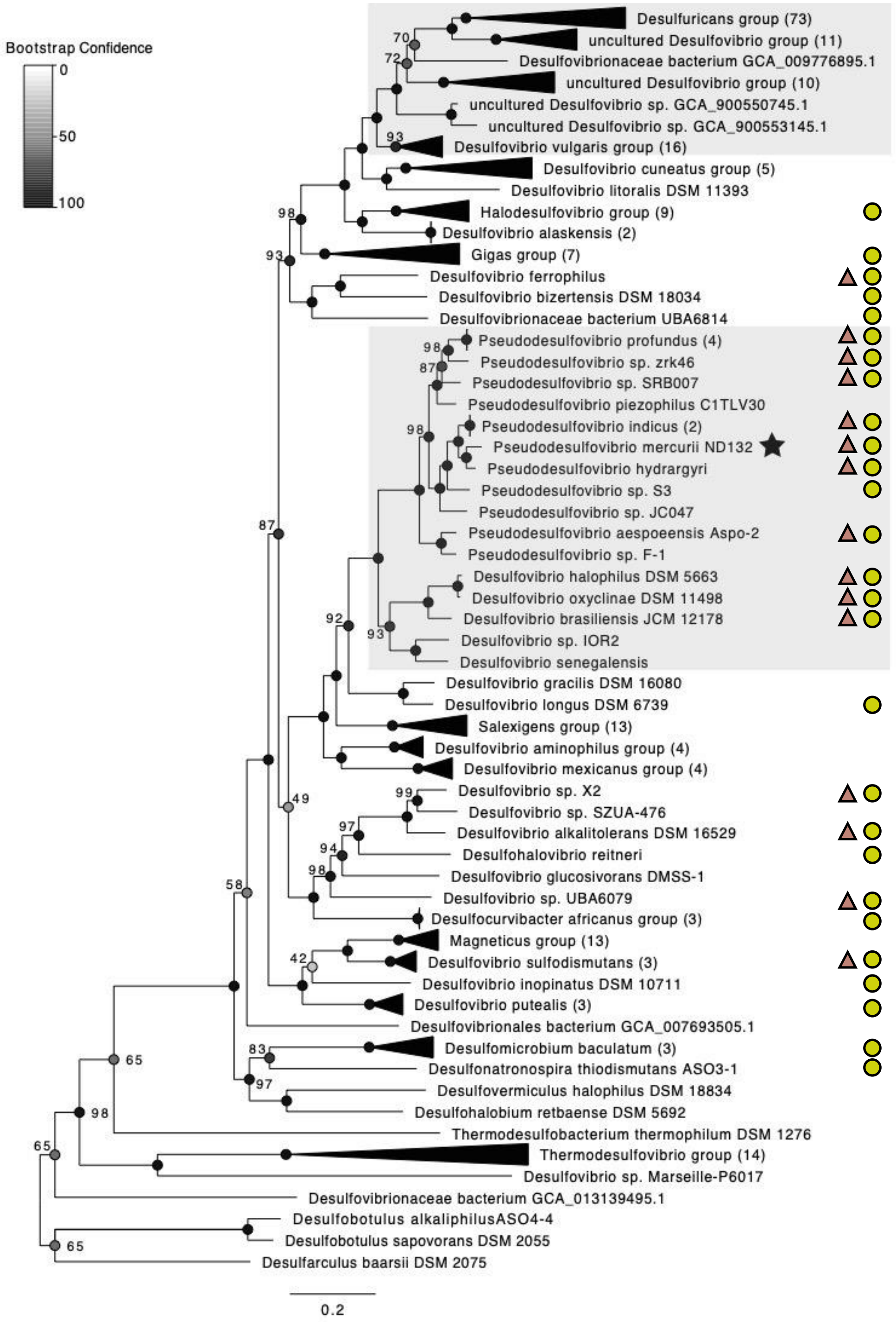


**Supplemental Figure S1**. Phylogenetic tree of *Desulfovibrio* and *Pseudodesulfovibrio* genomes closely related to *Pseudodesulfovibrio mercurii* ND132 adapted from Gilmour et al. 2021(7). Maximum-likelihood tree built from sequence alignment of reference marker protein sequences using CheckM and RAxML. Genomes that encode HgcAB indicated by yellow circle, genomes with ArsR-like transcriptional regulator encoded directly upstream of *hgcAB* shown by brown triangle.

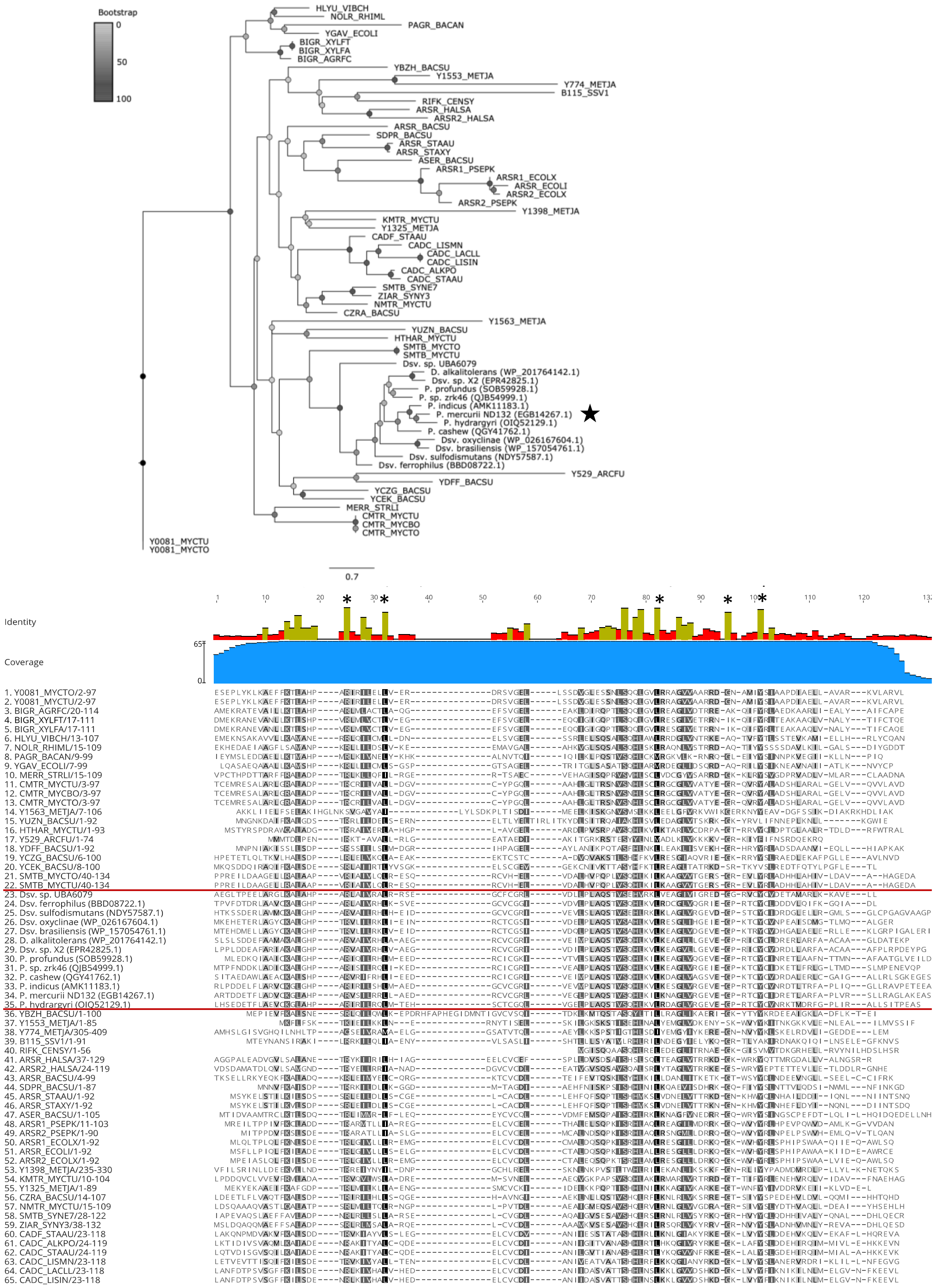


**Supplemental Figure S2**. Phylogenetic relationship between ArsR-like regulator (ArsR2, DND132_1054) in *Pseudodesulfovibrio mercurii* ND132 to other ArsR/SmtB-family transcriptional regulators. Reference sequences for the ArsR-type HTH domain profile were pulled from Expasy Prosite (Reference ID: PS50987). Included in red box are ND132 ArsR2 homologues from *Pseudodesulfovibro* and *Desulfovibrio* (Figure S2). Tree built using RAxML using LG+Gamma substitution model and 1000 rapid bootstraps from the protein alignment shown. The conserved invariant residues associated with the HTH family of regulators are denoted by an asterisk (*), the four-cysteine motif conserved in putative *hgcAB* regulators highlighted in yellow.

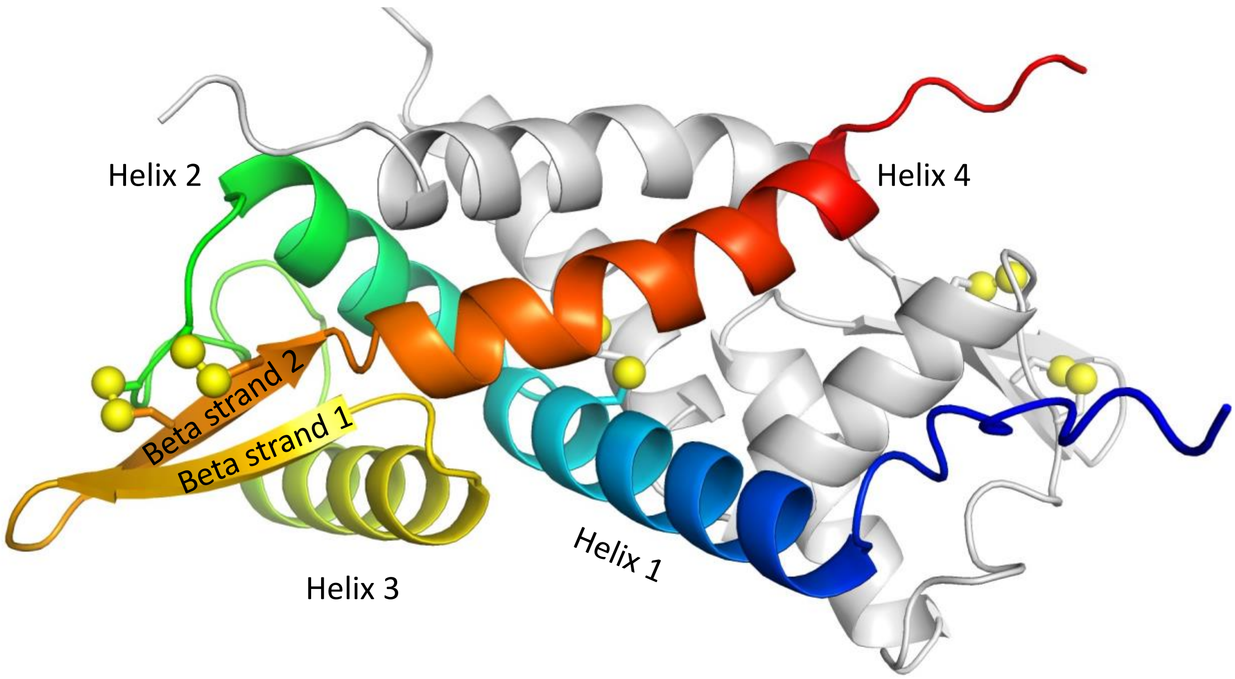


**Supplemental Figure S3.** Predicted metal(loid) binding site of the putative ArsR-like regulator from *P. mercurii* ND132 (DND132_1054). Chain A is shown in spectrum coloring (Blue = N-terminus, red = C-terminus) and chain B is in gray. Sulfur atoms of cysteine residues are shown as yellow spheres. Secondary structure elements are labeled.

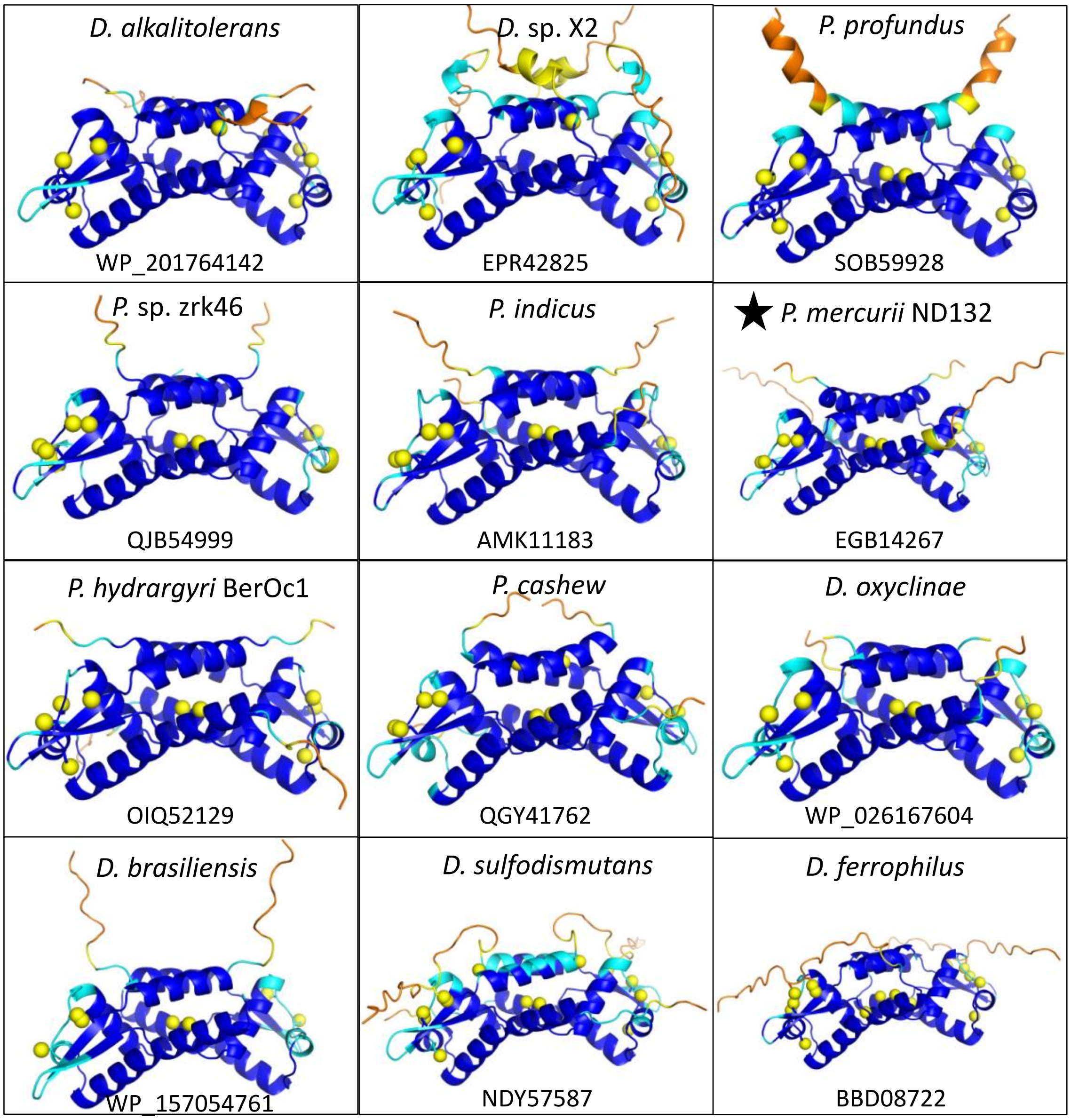


**Supplemental Figure S4**. AlphaFold2 models of selected ArsR-like proteins. Models are colored by predicted LDDT scores. Dark blue = very high confidence (pLDDT > 90), cyan = confident (90 > pLDDT > 70), yellow = low confidence (70 > pLDDT > 50), and orange = very low confidence (pLDDT < 50). Sulfur atoms of cysteine residues are shown as yellow spheres. *Desulfohalovibrio alkalitolerans* (WP_201764142.1), *Desulfovibrio* sp. X2 (EPR42825), *Pseudodesulfovibrio profundus* (SOB59928), *Pseudodesulfovibrio* sp. Zrk46 (QJB54999), *Pseudodesulfovibrio indicus* (AMK11183), *Pseudodesulfovibrio mercurii* ND132 (EGB14267), *Pseudodesulfovibrio hydrargyri* BerOc1 (OIQ52129), *Pseudodesulfovibrio cashew* (QGY41762), *Desulfovibrio oxyclinae* (WP_026167604), *Desulfovibrio brasiliensis* (WP_157054761), *Desulfovibrio sulfodismutans* (NDY57587), *Desulfovibrio ferrophilus* (BBD08722)

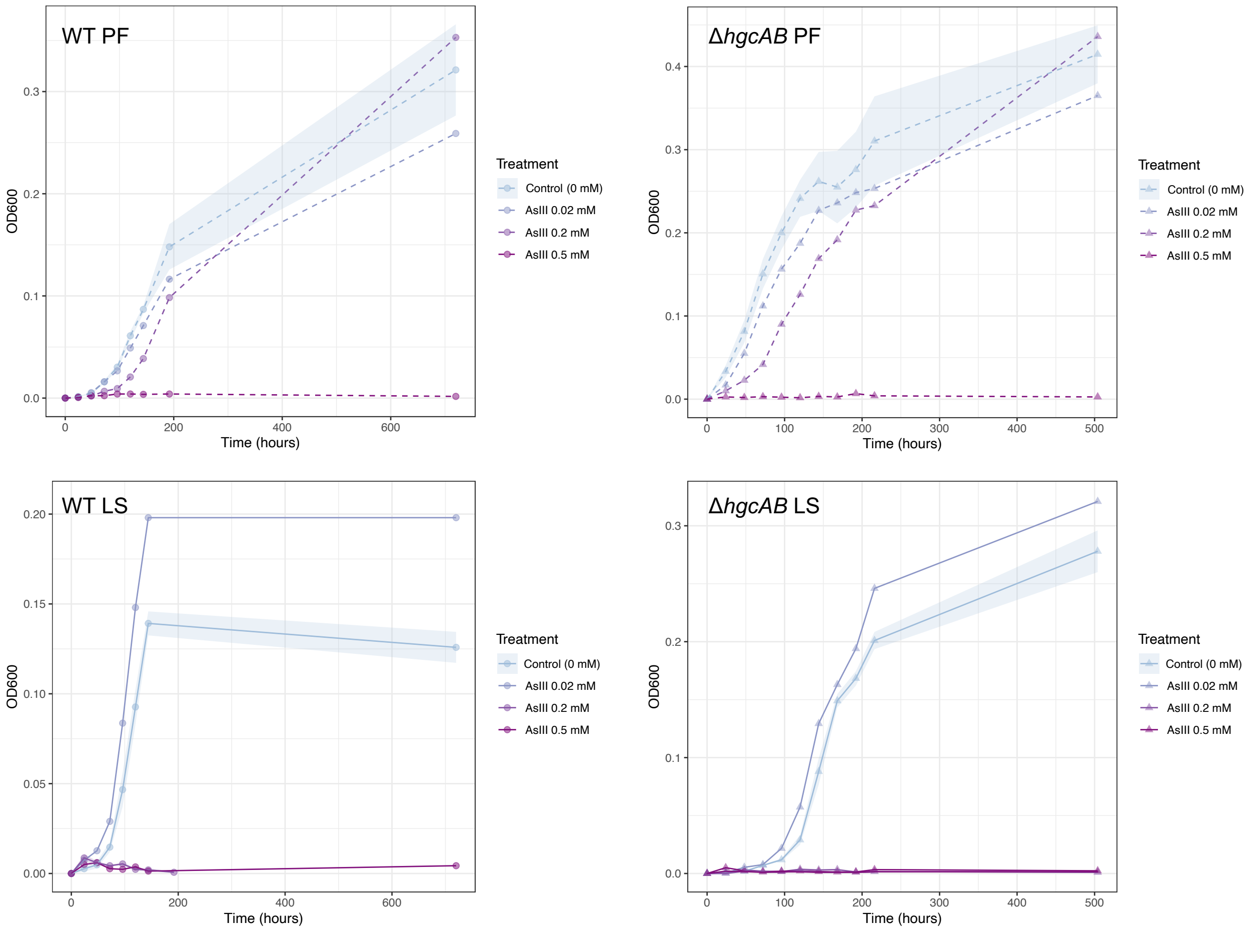


**Supplemental Figure S5**. Growth assays of tests of arsenite toxicity to *Pseudodesulfovibrio mercurii* ND132 wild-type (WT) and *hgcAB* deletion mutant (ΔhgcAB) in defined CCM media with pyruvate-fumarate (PF) or lactate-sulfate (LS) and sodium arsenite, As(III). Growth curves fitted to optical density measured at 600 nm absorbance of biological triplicates using 'ggplot' and 'dplyr' in R. Shaded area shows the standard error of average $OD_{600}$ measurements for control replicates.

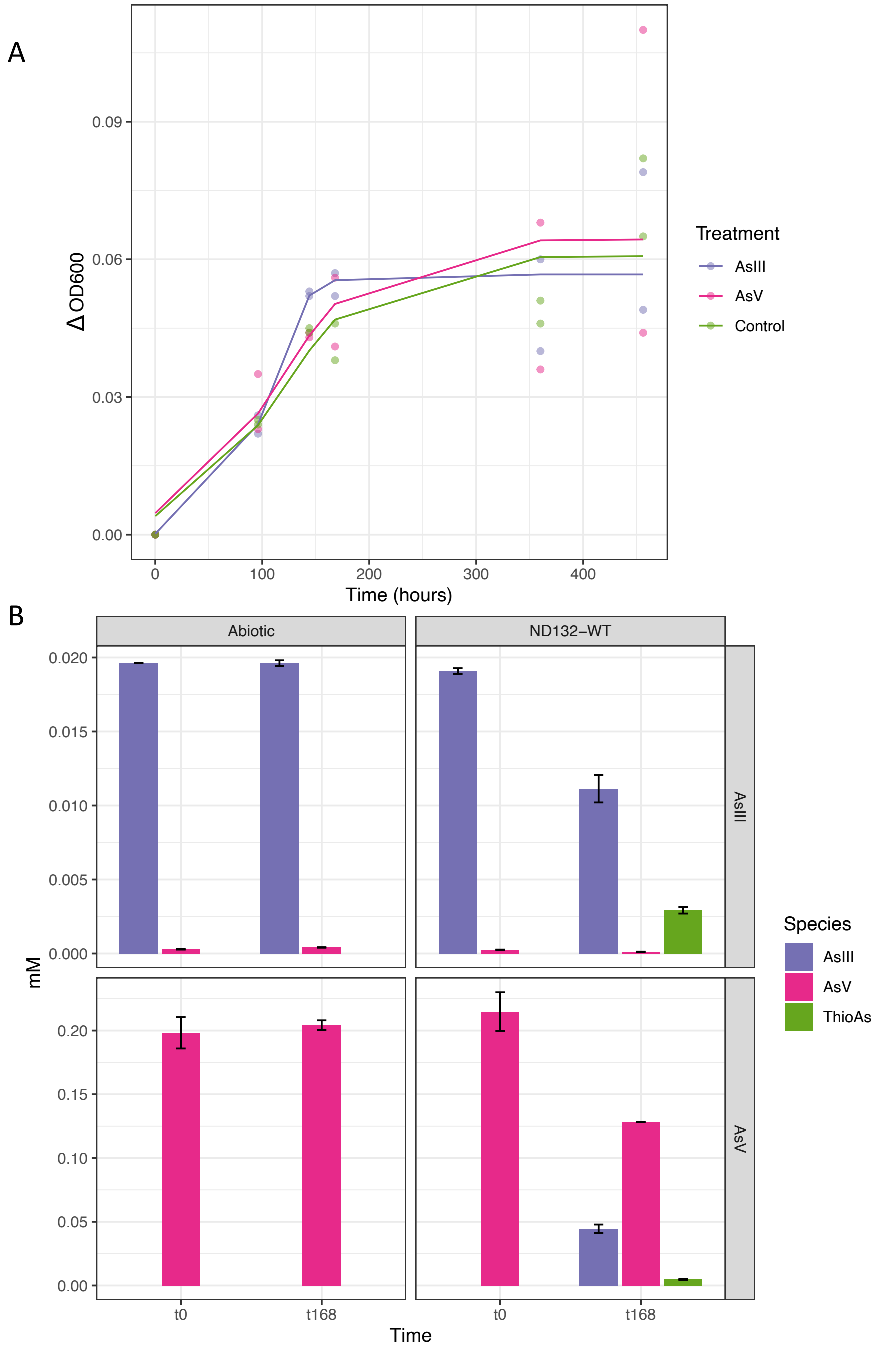


**Supplemental Figure S6**. (A) Growth curve of *Pseudodesulfovibrio mercurii* ND132 wild-type cultures in CCM-PF media with 0.02 mM sodium arsenite (AsIII), 0.2 mM sodium arsenate (AsV), or no arsenic (control). Growth curves fit to change in optical density at 600 nm absorbance (OD600) of biological duplicates using R program, Growthcurver(8). Culture media with arsenic was sampled before inoculation at t0 and after cells reached stationary phase, t = 168 hours for arsenic speciation analyses (B).

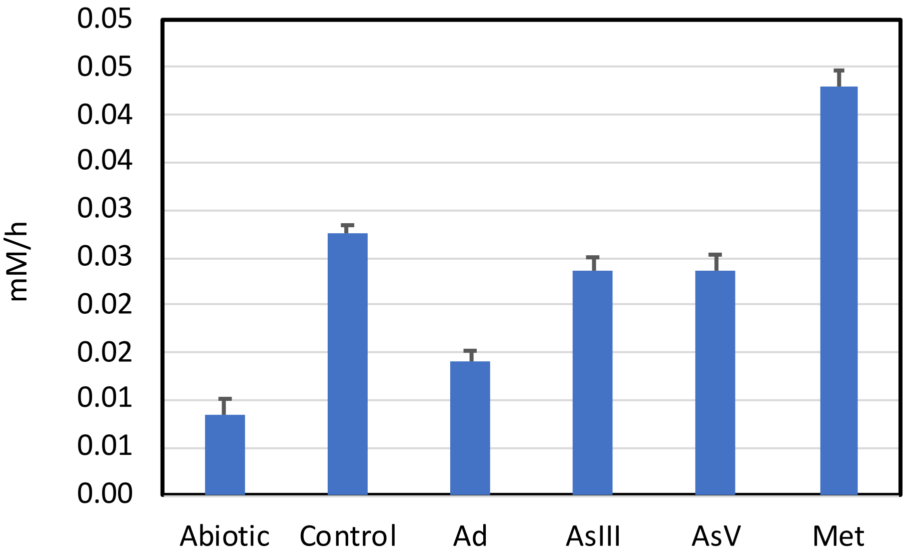


**Supplemental Figure S7**. Total inorganic carbon production (mM per hour) in the washed cell assays by treatment. Error bars are the standard error of the slope for duplicate assays. Based on paired analysis of the slopes of DIC+CO2 production, with α=0.050: Met>Control>AsIII=AsV>Ad>Abiotic

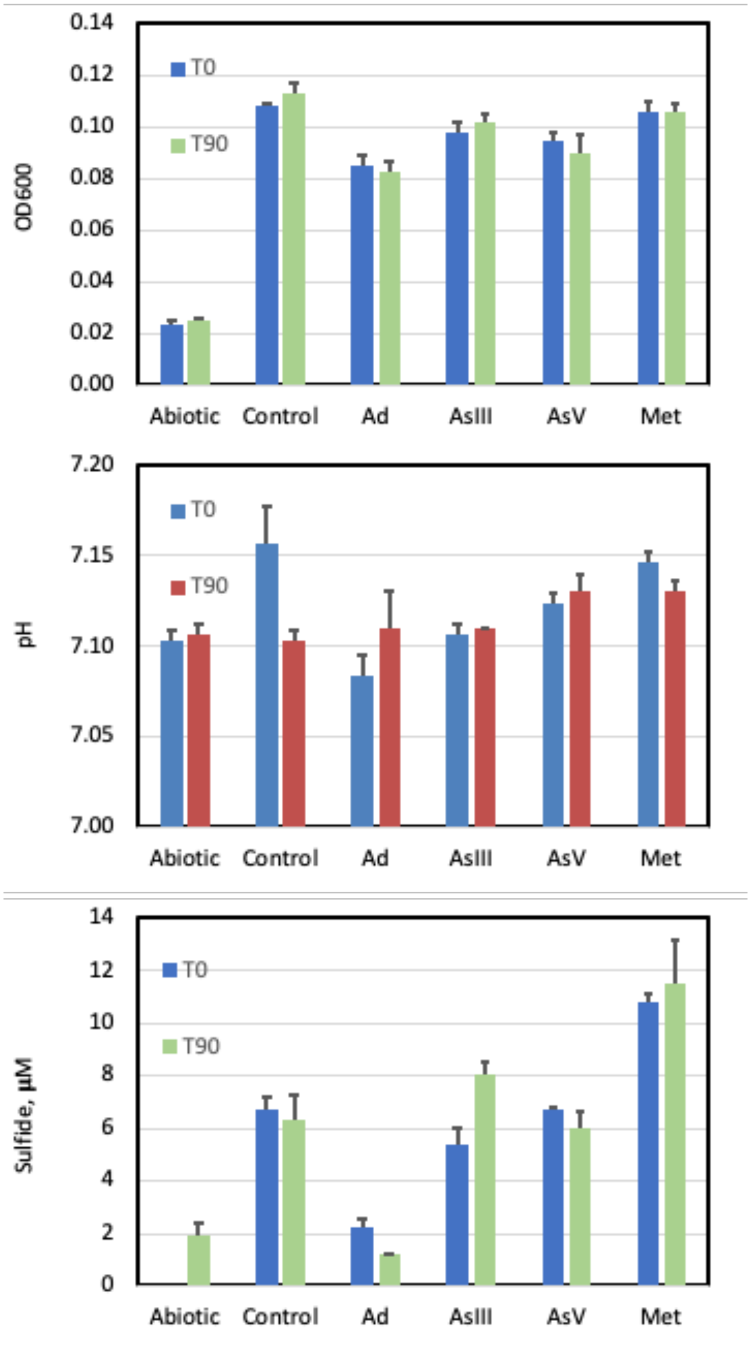


**Supplemental Figure S8**. Optical density (top), pH (middle) and sulfide (bottom) for washed cell assays at the beginning and end of methylation assays, by treatment. Error bars represent sStandard deviations from triplicate assays. For OD600, across both time points: Control=Met≥AsIII>AsV>Ad>Abiotic. For pH, there were no significant differences among treatments. For sulfide, across both time points: Met>AsIII=Control=AsV>Ad=Abtioic

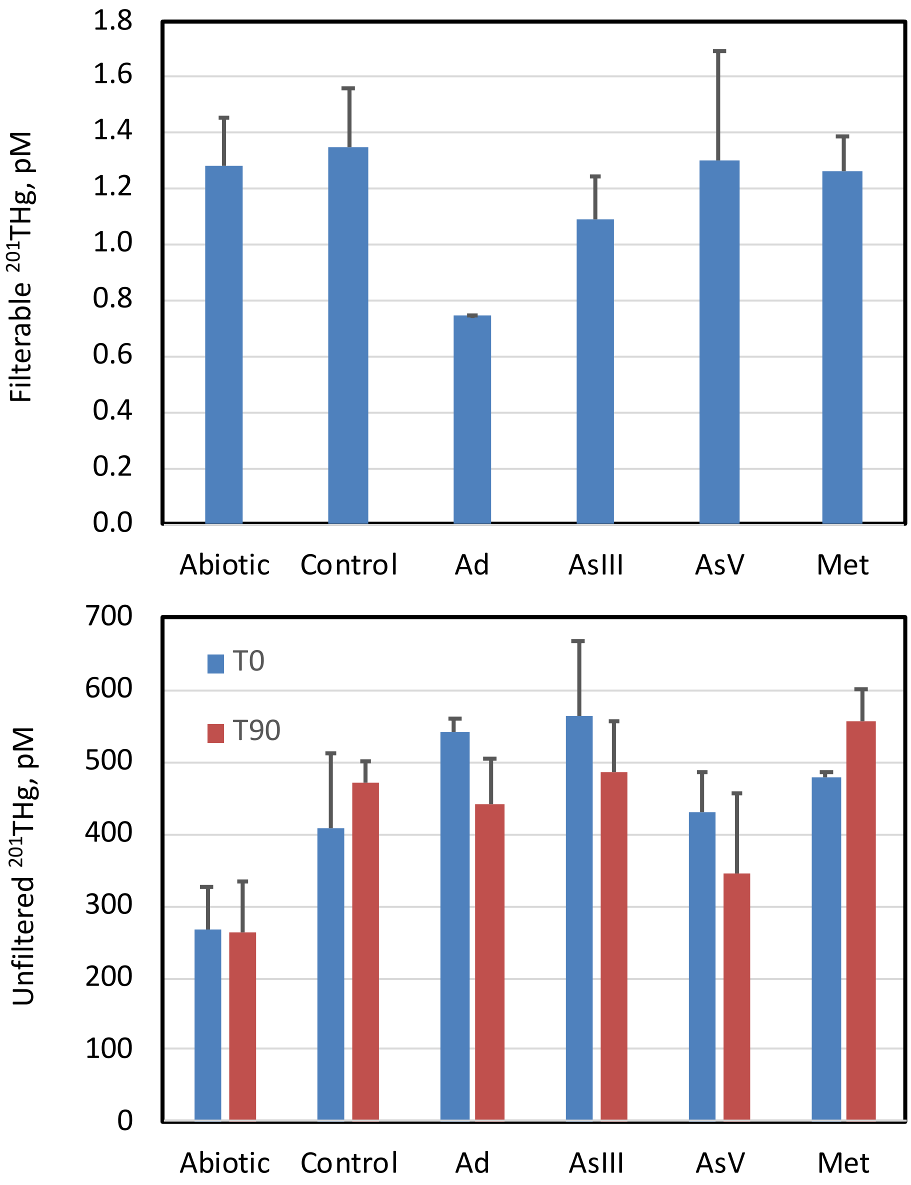


**Supplemental Figure S9**. Top, initial filterable $^{201}Hg$ concentrations in washed-cell assays, immediately after the addition of 1 nM $^{201}Hg$. Bottom, mercury mass balance in cultures, total unfiltered $^{201}Hg$ concentrations at the beginning and end of methylation assays.

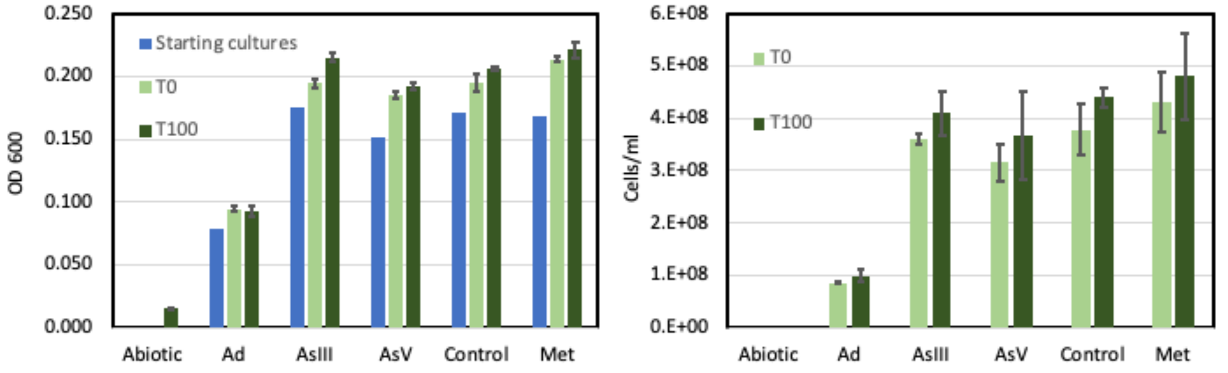


**Supplemental Figure S10**. Optical and cell density for ND132 batch cultures at the beginning and end of methylation assays, by treatment. Optical and cell density rose significantly during the assays. Error bars represent sStandard deviations from triplicate assays. For optical density: Met=AsIII=Control≥AsV>Ad. For cell counts: Met>AsIII=Control≥AsV>Ad. Both based on a Tukey least square means test of T0 and T100 data, with α=0.050.

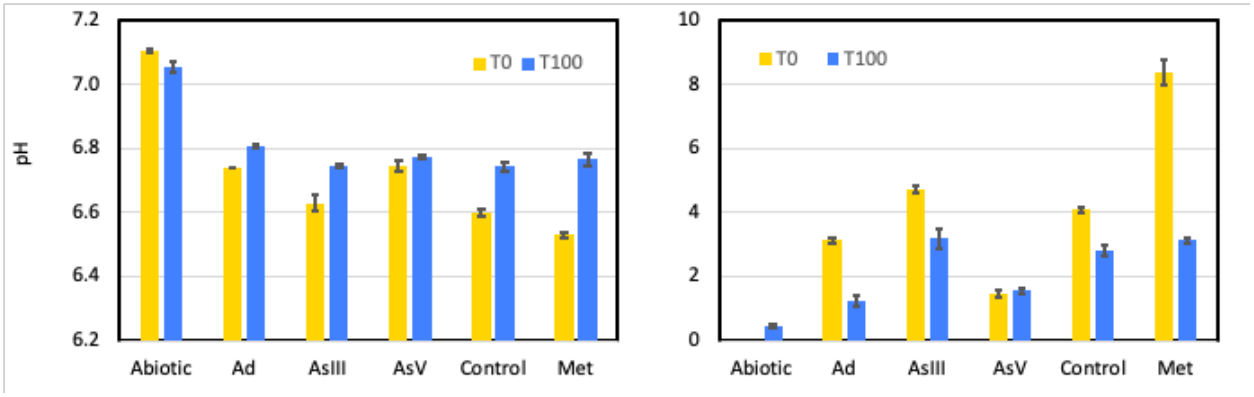


**Supplemental Figure S11**. Medium chemistry at the beginning and end of batch culture methylation assays, by treatment. Error bars represent s~~S~~tandard deviations from triplicate assays. Sulfide declined and pH rose significantly during the assays. For pH: Abiotic>Ad=AsV>AsIII=Control≥Met. For sulfide: Met>AsIII>Control>Ad>AsV>Abiotic. Both based on a Tukey least square means test of T0 and T100 data, with α=0.050.

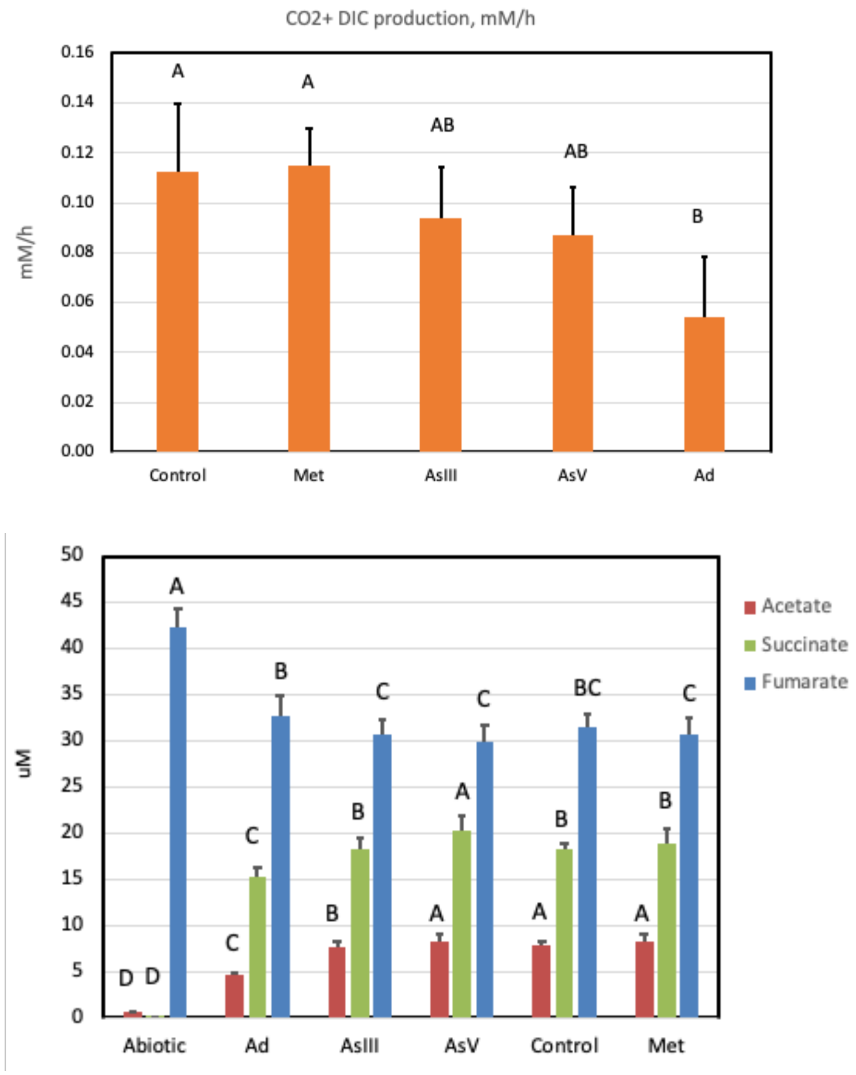


**Supplemental Figure S12**. Metabolic activity of treated *Pseudodesulfovibrio mercurii* ND132 cultures as assayed by (top) total dissolved inorganic carbon (DIC) production and organic acid concentrations (bottom). For DIC production, data are the average of two biological replicate assays, error bars are the standard error of the slopes. Differences are based on pairwise comparison of slopes ($p<0.050$). For organic acids, error bars are standard deviations from triplicate assays. Levels not connected by same letter are significantly different based on a Tukey least square means test with $\alpha=0.050$.

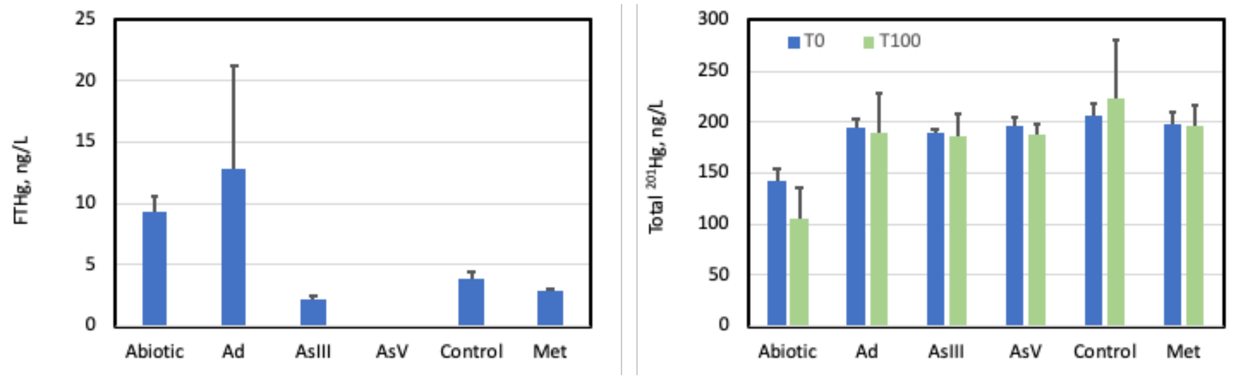


**Supplemental Figure S13**. Left, initial filterable $^{201}Hg$ concentrations in cultures, immediately after the addition of 1 nM $^{201}Hg$. Right, mercury mass balance in cultures, total unfiltered $^{201}Hg$ concentrations at the beginning and end of methylation assays. Error bars are standard deviations for triplicate assays.

A

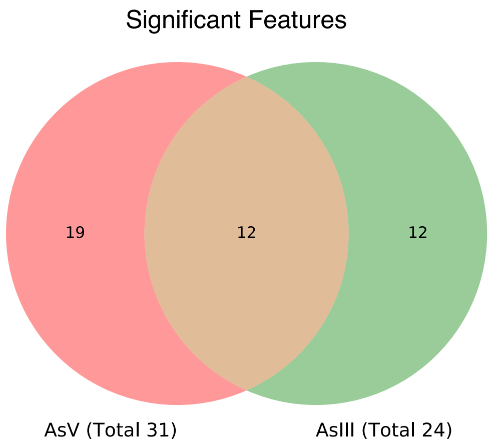


B

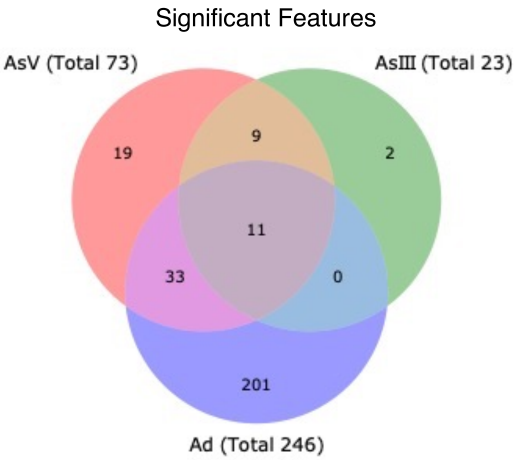


**Supplemental Figure S14**. Venn diagrams showing shared features with significant (|logFC| > 1.5, $p < 0.05$) differential expression in RNA-seq datasets from (A) washed-cell and (B) batch culture assays. Differential expression in treatment vs. control, with only AsIII and AsV being shown for washed-cells and AsIII, AsV, and adenosine (Ad) for batch cultures.

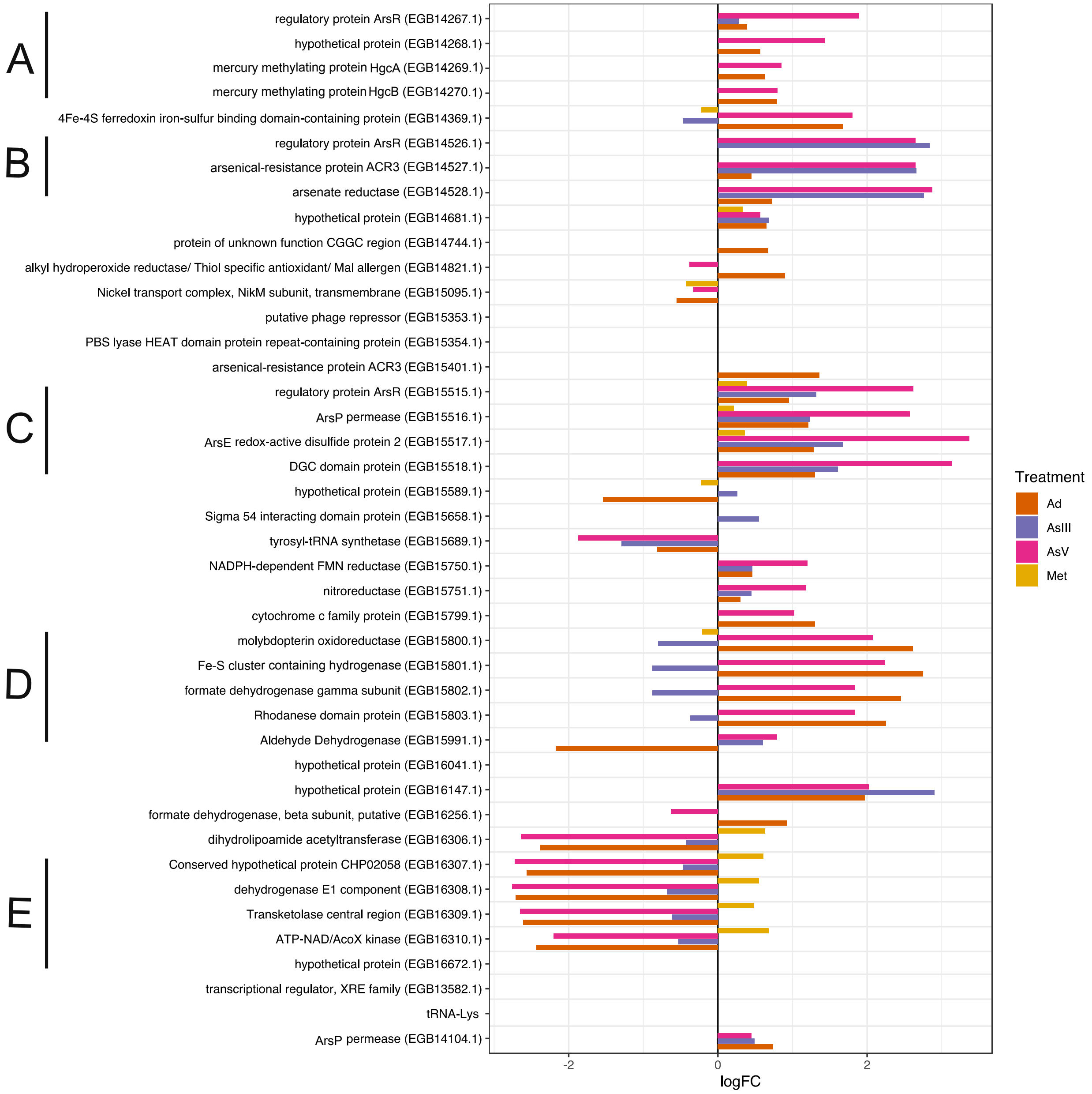


**Supplemental Figure S15**. Differential expression of significant features ($p < 0.05$) from RNA-seq analyses of adenosine dialdehyde (Ad, 0.1 mM), methionine (Met, 1 mM), As(V) (0.2 mM) and As(III) (0.02 mM) treated batch cultures from Hg methylation assays of wild-type *Pseudodesulfovibrio mercurii* ND132. These correspond to the significant features that were differentially expressed in the transcriptomes of arsenic treated washed-cells (Figure 6), including several putative operons (A-E).

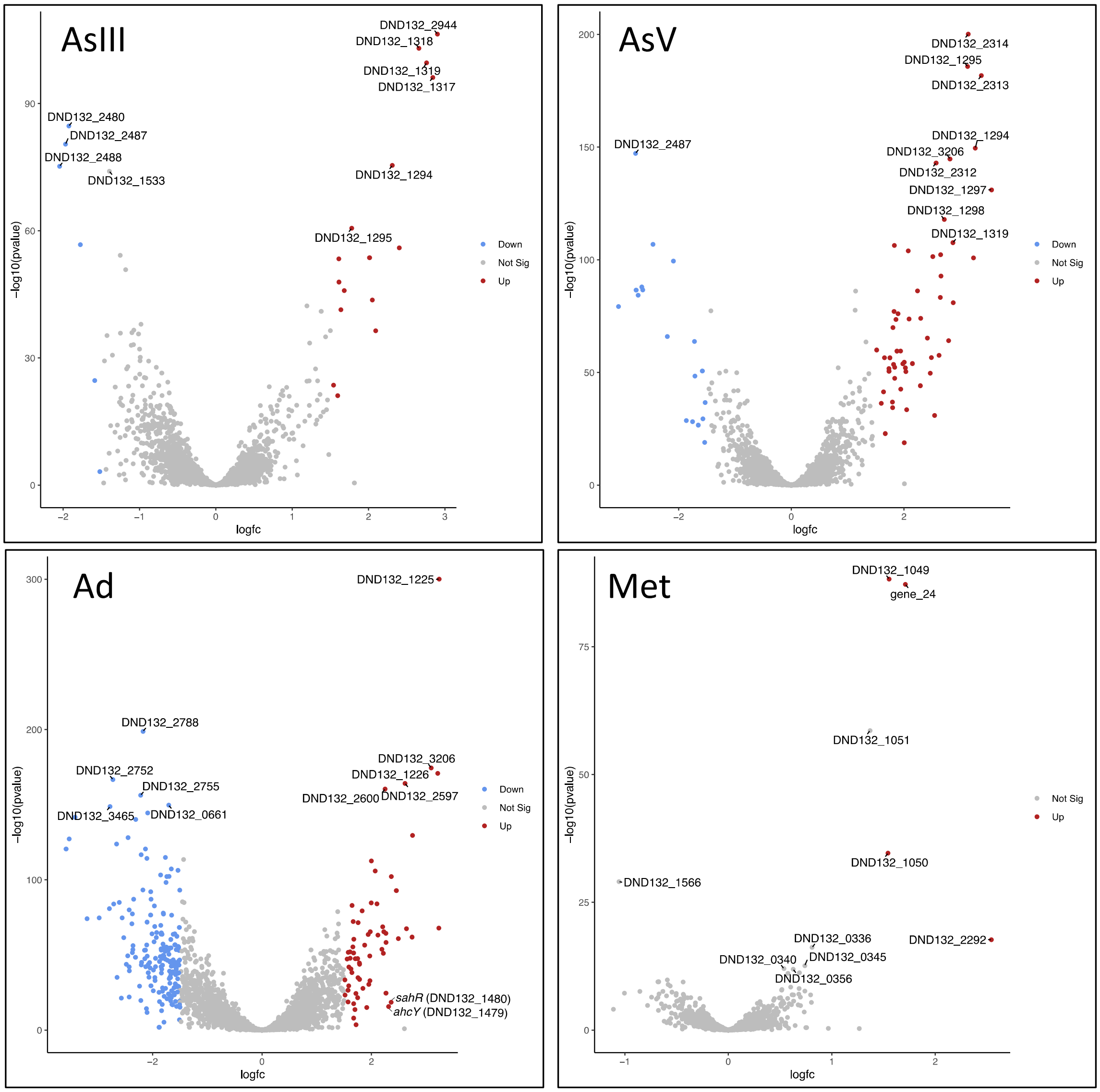


**Supplemental Figure S16**. Volcano plots showing differential expression of features from batch culture RNA-seq datasets. Red dots indicate significantly up-regulated features (logFC > 1.5, $p < 0.05$), while blue indicates significantly down-regulated features (logFC < -1.5, $p < 0.05$). The top ten most significant differentially expressed features in each RNA-seq dataset are labeled by gene ID.

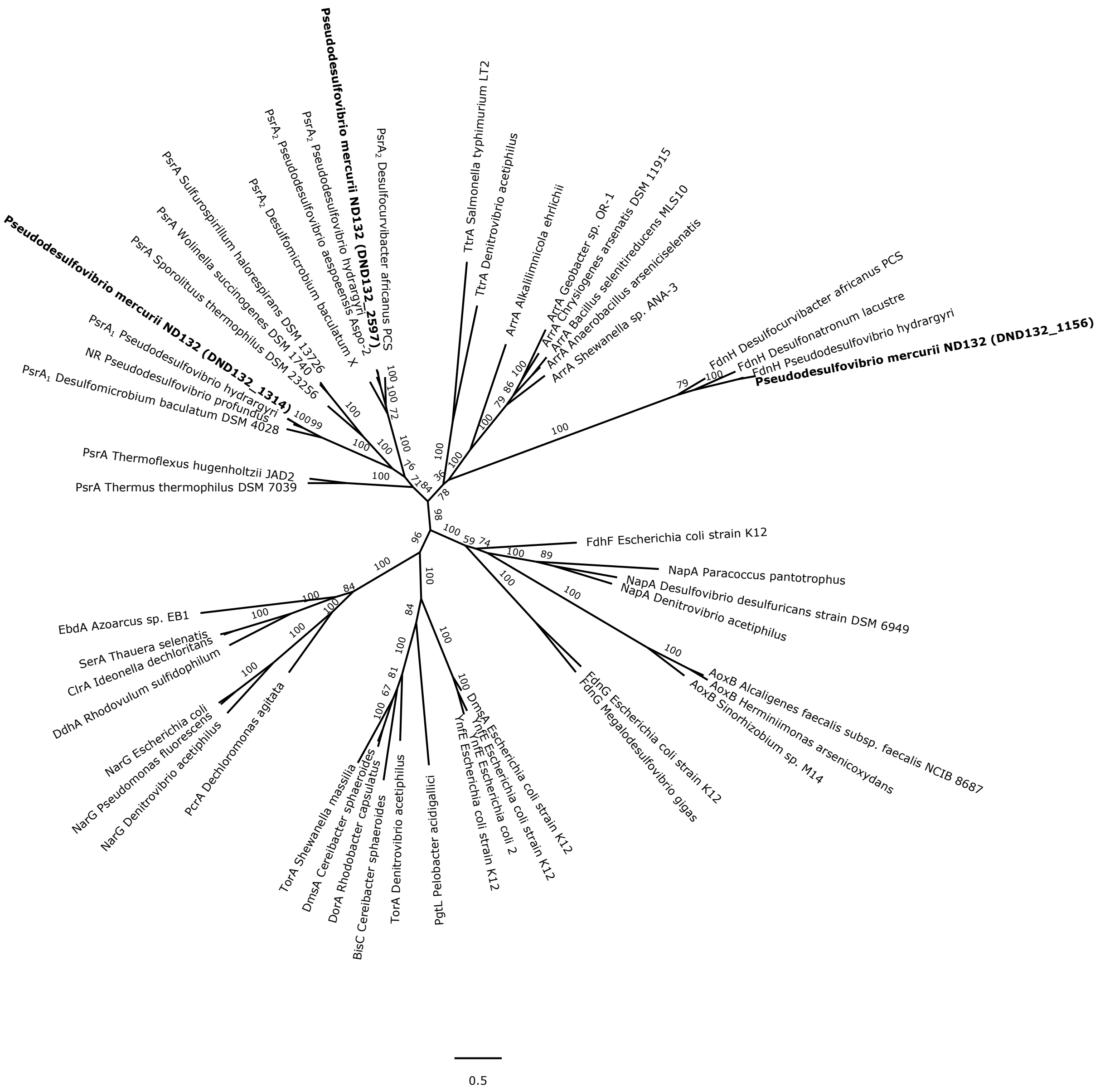


**Supplemental Figure S17**. Phylogenetic relationship of the DMSO-molybdopterin family of proteins. The three protein sequences identified in the *Pseudodesulfovibrio mercurii* ND132 genome are shown in bold. The unrooted tree was constructed using maximum likelihood analysis in RAxML, and bootstrap values were calculated by 100 resamplings. ArrA, arsenate reductase; AoxB, arsenite oxidase; BisC, biotin sulfoxide reductase; ClrA, chlorate reductase; DmsA/Dor/Tor/YnfE, dimethyl sulfoxide/ trimethylamine N-oxide reductase; ; Fdh, formate dehydrogenase; Nar/NR, nitrate reductase; PadB, phenylacetyl-CoA dehydratase; PsrA/PhsA, polysulfide/polythionate reductase; SerA, selenate reductase; TorA, trimethylamine N-oxide reductase; TtrA, tetrathionate reductase.

**Supplemental Materials References**


1. Horvat M, Liang L, Bloom NS. 1993. Comparison of distillation with other current isolation methods for the determination of methyl mercury compounds in low level environmental samples: Part II. Water. Analytica Chimica Acta 282:153-168.
2. De Wild JF, Olsen ML, Olund SD. 2002. Determination of methyl mercury by aqueous phase ethylation, followed by gas chromatographic separation with cold vapor atomic fluorescence detection. Survey USG,
3. Tutschku S, Schantz MM, Horvat M, Logar M, Akagi H, Emons H, Levenson M, Wise SA. 2001. Certification of the methylmercury content in SRM 2977 Mussel Tissue (organic contaminants and trace elements) and SRM 1566b Oyster Tissue. Fresenius' Journal of Analytical Chemistry 369:364-369.
4. Edgar RC. 2004. MUSCLE: multiple sequence alignment with high accuracy and high throughput. Nuc Acids Res 32:1792-1797.
5. Capo E, Peterson BD, Kim M, Jones DS, Acinas SG, Amyot M, Bertilsson S, Björn E, Buck M, Cosio C. 2022. A consensus protocol for the recovery of mercury methylation genes from metagenomes. Molecular Ecology Resources.
6. Gionfriddo C, Capo E, Peterson B, Lin H, Jones D, Bravo A. 2021. Hg-cycling Microorganisms in Aquatic and Terrestrial Ecosystems Database v1. 01142021.
7. Gilmour CC, Soren AB, Gionfriddo CM, Podar M, Wall JD, Brown SD, Michener JK, Urriza MSG, Elias DA. 2021. Pseudodesulfovibrio mercurii sp. nov., a mercury-methylating bacterium isolated from sediment. International Journal of Systematic and Evolutionary Microbiology 71:004697.

8. Sprouffske K, Wagner A. 2016. Growthcurver: an R package for obtaining interpretable metrics from microbial growth curves. BMC Bioinformatics 17:172.
